# Quantification of Signal Amplification for Receptors: The *K*_d_/EC_50_ Ratio of Full Agonists as a Gain Parameter

**DOI:** 10.1101/2024.10.17.618878

**Authors:** Peter Buchwald

## Abstract

Concentration-response relationships connecting the concentration of ligands to the responses they produce are central to pharmacology in general and form the core of quantitative pharmacology. While typically they can be well-described by hyperbolic functions (sigmoid on commonly used semi-log scales) and characterized by half-maximal concentrations values (EC_50_), their connection to receptor occupancy, characterized in a similar manner by the equilibrium dissociation constant *K*_d_, can be complex due to the intermixing of the effects from occupancy-induced activation with those from partial agonism, constitutive activity, and pathway-specific signal amplification. Here, it is proposed that, as long as both occupancy and response follow such typical concentration-dependencies, signal amplification can be quantified using the gain parameter *g_K_*=*κ*=*K*_d_/EC_50_ measured for full agonists. This is similar to the gain parameter used in electronics (e.g., *g_V_*=*V*_out_/*V*_in_ for voltage). On customarily used semi-log representations, log *g_K_* corresponds to the horizontal shift between the response and occupancy curves, log*K*_d_-logEC_50_, the presence of which (i.e., *K*_d_>EC_50_) is generally considered as evidence for the existence of “*receptor reserve*” or “*spare receptors*”. The latter is a misnomer that should be avoided since even if there are excess receptors, there is no special pool of receptors “not required for ordinary use” as *spare* would imply. For partial agonists, the *κ*=*K*_d_/EC_50_ shift is smaller than for full agonists as not all occupied receptors are active. The *g_K_* gain parameter (full agonist *K*_d_/EC_50_) correspond to the *γ* gain parameter of the SABRE receptor model; for partial agonists (*ε*<1), SABRE predicts a corresponding shift of *κ*=*εγ*-*ε*+1.

## Introduction

### Receptors and Concentration- or Dose-Response Curves

Receptors, “pharmacology’s big idea” [1], are at the core of our current understanding of mechanism of drug action (Figure 1) [2-5]. In a general context, receptors are used to denote any target of a substance that is responsible for initiating a biological response, but in a stricter pharmacological sense, receptors are protein molecules whose function is to recognize and respond to endogenous chemical signals; as such, they are one of the four common drug targets (receptors, enzymes, carriers, and ion channels) [6]. Binding of a ligand to the receptor initiates a sequential process termed signal transduction that culminates in one or more specific cellular response. Unequivocally connecting the concentration of a ligand of interest to the response it produces is of obvious interest; thus, concentration-response relationships are central to pharmacology in general and quantitative pharmacology in particular. Typically, the concentration-dependence of both receptor response and occupancy can be well described by hyperbolic functions (sigmoid on commonly used semi-log scales) characterized by half-maximal concentrations values, i.e., *K*_obs_ = EC_50_ and *K*_d_ for response and occupancy, respectively (Figure 2). These are determined from single-parameter equations such as those shown below for fractional occupancy (*f*_occup_) and response (*f*_resp_), respectively:

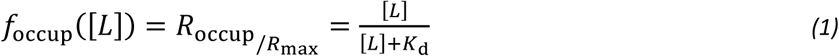

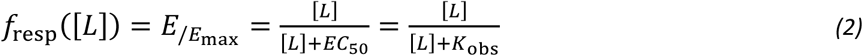

**Figure 1.**
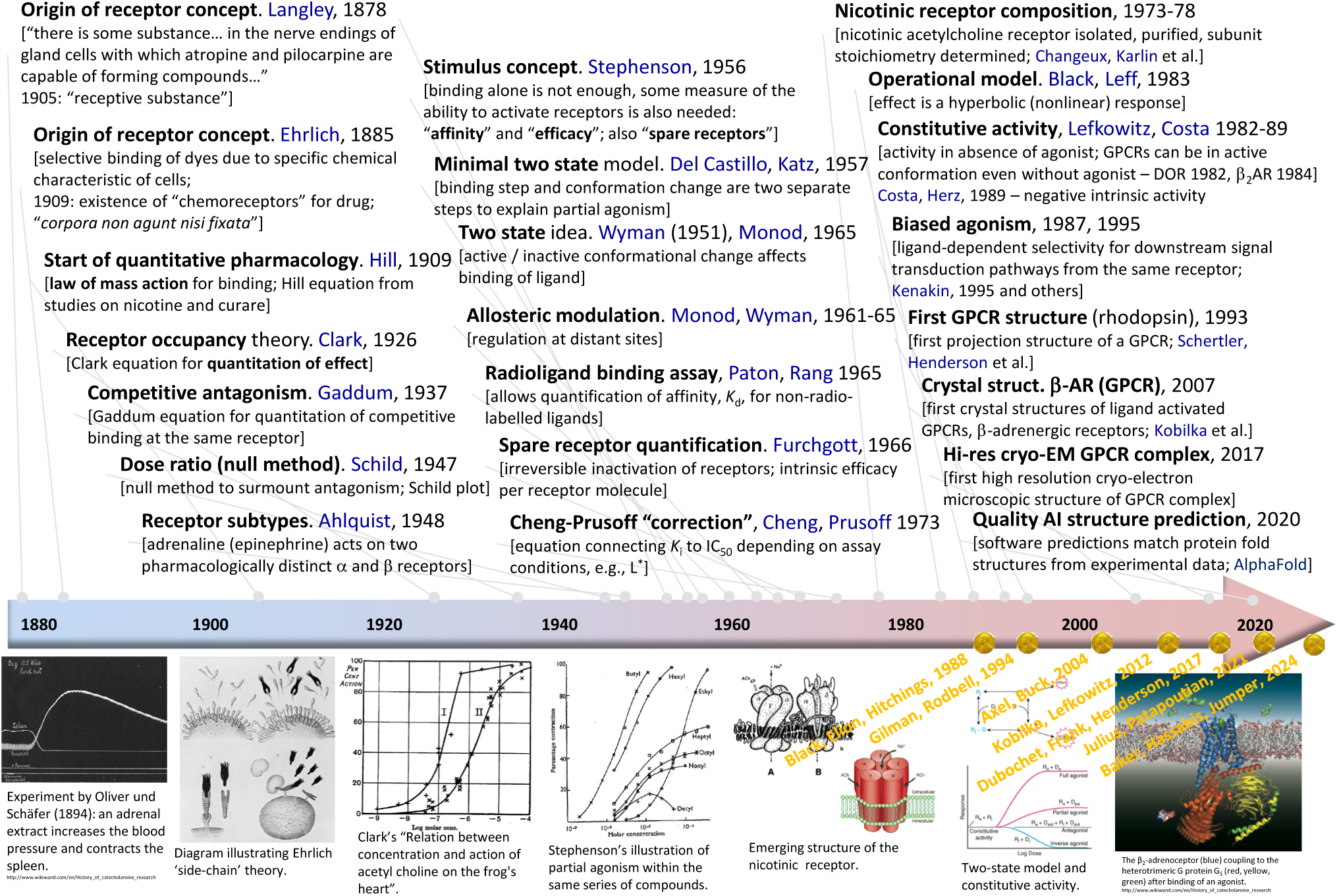
Timeline highlighting some of the main receptor-related developments. For each entry, the main idea, the year of its introduction, and the names of the people involved are listed. A few representative illustrations are included in the bottom row in chronological order; names and years in yellow shown there highlight Nobel prizes awarded for what can be considered as receptor-related discoveries.

**Figure 2.**
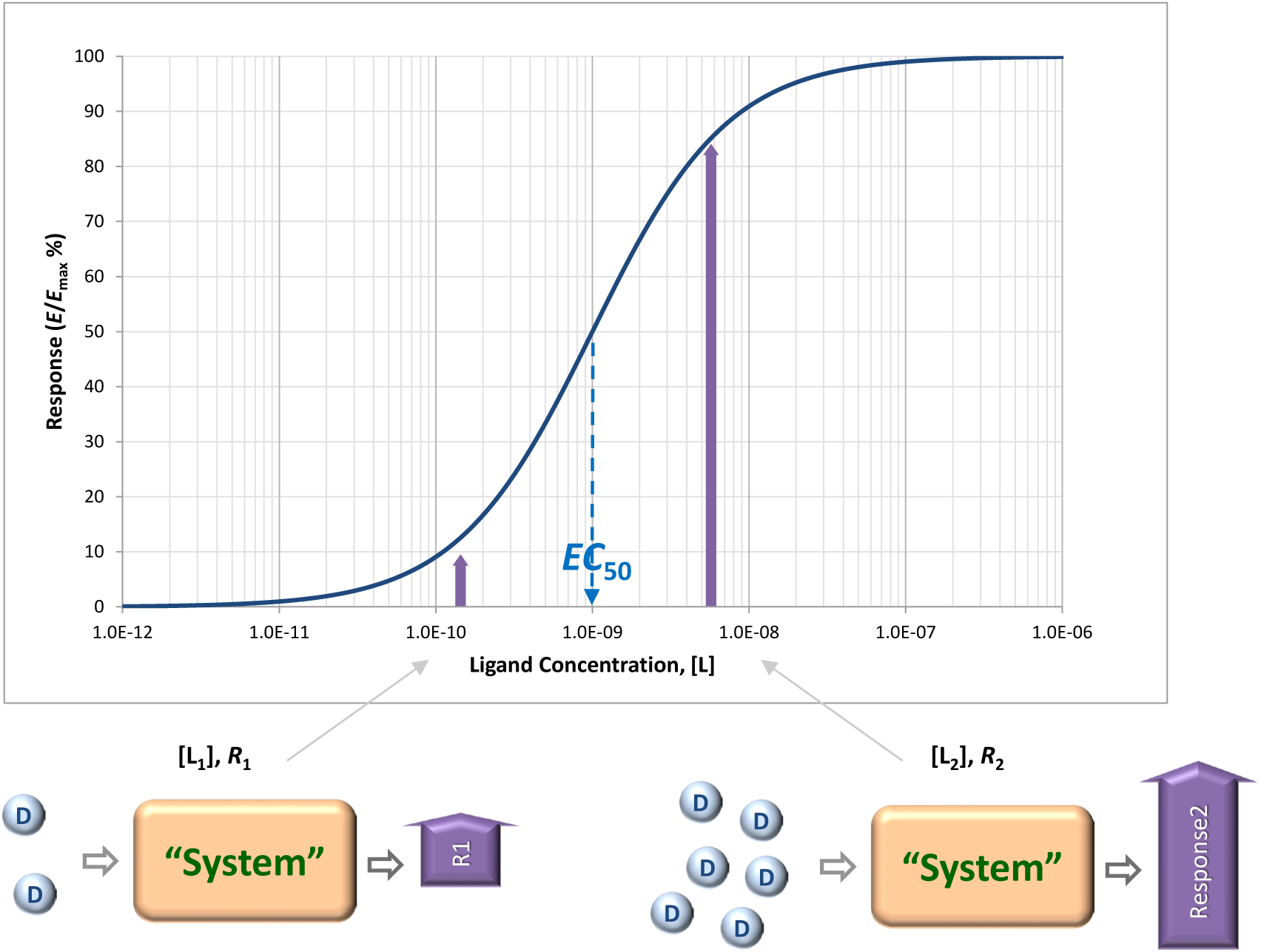
Concentration-response relationships connecting the concentration of ligands to the responses they produce are central to pharmacology. Concentration-response curves, and even dose-response curves obtained in *in vivo* systems, are typically well described by hyperbolic functions (sigmoid on commonly used semi-log scales as shown here) and characterized by half-maximal concentrations values, EC_50_ (eq. 2).

With the evolution of the field (see summary timeline of Figure 1), it became clear that the connection between receptor occupancy and response can be quite complex, as in addition to occupancy, it also has to account for partial agonism, receptor reserve, constitutive activity, and other effects. The overall efficiency of the transduction process that links occupancy to response, i.e., the occupancy–response coupling [7], is influenced by what takes place at the receptor (i.e., how many receptors are occupied by the ligand, what fractions of the occupied and unoccupied receptors are active) as well as by what takes place “downstream” from the receptor (i.e., the biochemical events that transduce the “active” signal from the receptor into the response of interest). Thus, the generated response depends not only on • occupancy, which is determined by *ligand affinity* (i.e., the ability of the ligand to bind to the receptor), but also on • the ability of the ligand to activate the receptor upon binding, which is determined by the *efficacy* of the ligand, as there are partial agonists that cannot fully activate occupied receptors, • the degree of the possible activation of unoccupied receptors, which should be quantifiable via an *efficacy of the constitutive activity* of the receptor, as there are constitutively active receptors, • the pathway-dependent *signal amplification*, which should be quantifiable by a *gain parameter*, as responses can run “ahead” of occupancy, i.e., are more sensitive to concentration than occupancy so that concentration-response curves are left-shifted compared to concentration-occupancy curves, and • the steepness of the concentration dependence, which is characterizable by a *Hill slope* or coefficient, as concentration response can be more or less abrupt than those strictly following the law of mass action [8]. Hence, even if the simple Clark equation (eq. 2), which has been published a century ago in 1926 (Figure 1) [9]^1^, works well and forms the basis of most concentration-response or dose-response relationship fittings to this day, a quantitative model that can form the basis of a true concentration-response relationships and connect the concentration of a ligand to the response(s) that it produces should account for all these and, thus, have a minimum of five independent parameters [8]. Most likely, even more parameters are needed for dose-response relationships (*in vivo* systems), where the pharmacokinetic aspects determining the amount (concentration) of drug ultimately reaching the receptor also have to be accounted for [11, 12].

### Quantifying Signal Amplification for Receptors

Along these lines, it has been clearly recognized since the mid-1950s (Figure 1) that, for some receptors, maximal or close to maximal response can be achieved when only a much smaller fraction of the receptors is occupied – a recognition that led to the notion of “spare receptors” or “receptor reserve” [13, 14]. A method to quantify the fraction of occupied receptors (occupancy) from measurements of response (effect) data alone was introduced by Furchgott about a decade later [15, 16], and versions of it are still in use [17]. Quantitative studies comparing occupancy and response have been done for a number of cases, most of them involving G-protein coupled receptors (GPCRs), and some found quite extreme cases of activation. For example, guinea pig ileal response, where histamine can produce close to 100% response at only ∼2% occupancy (*f*_resp_ ≈ 1.00 at *f*_occup_ = 0.02), or rat heart β-adrenergic receptors, where epinephrine can produce half-maximal increase of muscle contractility at only ∼2% occupancy (*f*_resp_ ≈ 0.50 at *f*_occup_ = 0.02) (see [18] for additional references). This implies that significant signal amplification has to take place downstream – typically via some cascade mechanism involving second messengers such as cyclic AMP (cAMP) or other systems [19-21]; a classic textbook illustration for GPCRs can be found [here](https://open.lib.umn.edu/app/uploads/sites/211/2018/05/36.png) [22]. Sensory systems in particular need to rely on high gain, low noise amplification processes to be able to convert even very weak stimuli into detectable and reliable signals [23]. Along these lines, in phototransduction a single photon has been shown to activate around 60 G proteins from rod photoreceptors in frogs, which can lead to hydrolysis of up to 70,000 cGMP molecules downstream [24]. Consequently, humans seem to be able to detect a single-photon incident on the cornea [25]. Another well-known example is the mechanism by which epinephrine (adrenaline) or glucagon sets off a cascade of phosphorylation leading to the production of glucose resulting in a very strong amplification of the initial signal, possibly as high as 10^7^–10^8^-fold as (sub)nanomolar changes (<10^-9^ M) in these signals can lead to millimolar changes (∼10^-3^ M) in glucose [26, 27]. Signal amplification has been confirmed not only for GPCRs, but also for kinase-linked receptors [4] and ligand-gated ion channels (for example, for insect olfactory receptors whereby receptor excitation allows for direct influx of Ca^2+^ serving as second messenger and thereby amplify responses [28]). It is therefore important to have a well-defined method and corresponding parameter(s) that can be used to quantify receptor signal amplification.

## Methods

### Gain Quantification Based on Analogy with Electronic Amplifiers

Since the cascade mechanisms of physiological signal amplification resembles in many ways those occurring in electronic circuits [29], it makes sense to define a signal amplification factor in a manner similar to that used there. In a very general perspective, for a transfer function, *y* = *f*_tr_(*x*) that links the magnitude of the output response *y* to the input signal *x*, the gain (or amplification factor) *g* is defined as the ratio of output to input at a given value of input *x* [30]:

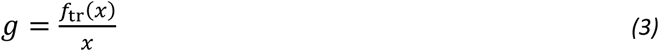

The input and output signals have to be of the same type (i.e., measured in the same units); therefore, the gain for voltage as a common example from electronics is:

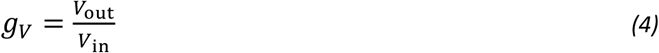

For linear amplifiers, the simplest case, the gain is constant over the entire range of the input, but in general, the value of the gain *g* depends on the input level, *x*. In such cases, the derivative of the transfer curve, d*f*_tr_/d*x*, is of interest as it represents the change in output (d*y*) produced by a small change in input (d*x*) at level *x* [30]. For linear amplifiers, this is identical to the gain *g* itself, as in this case, *f*_tr_(*x*) = *gx* for all *x* values with *g* being constant, so that d*f*_tr_/d*x* = *g*.

In physiological / pharmacological signaling, there can be significant increase in the number (concentration) of molecules due to downstream amplification from possibly only a few ligands that bind to the receptor (“input”) to a much larger number of generated molecules where the response or readout takes place (“output”) – a few well-known illustrative cases, such as that of epinephrine → glucose, have been mentioned earlier. The corresponding increase ratio could be used to calculate a concentration gain, *g_C_* = *C*_out_/*C*_in_; however, concentrations are rarely measured (or even measurable) along signaling pathways. On the other hand, responses are commonly measured (often normalized to their maximum) and quantified using EC_50_, the ligand concentration that causes half-maximal response (*f*_resp_ = 0.5; eq. 2). Thus, it makes sense to quantify amplification using them, especially as they are also measured in concentration units. One way to do so is to consider the signal that is amplified to be not the ligand concentration, but the effectiveness of the ligand to generate a response, i.e., the signaling power per unit input (molecule). Since potency quantification is done by respectively), one can consider the respective potency per ligand as proportional with the inverse of them. Thus, the signal per ligand is inversely proportional with the concentration of ligands needed to achieve a given, e.g., half-maximal effect, *σ_K_* = 1/*K*, and then the corresponding gain *g_K_* is the ratio of these, i.e., *σ_K_* at the level of occupancy as the “in” signal and *σ_K_* at the level of measured response as the “out” signal (Figure 3):

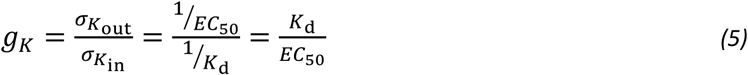

**Figure 3.**
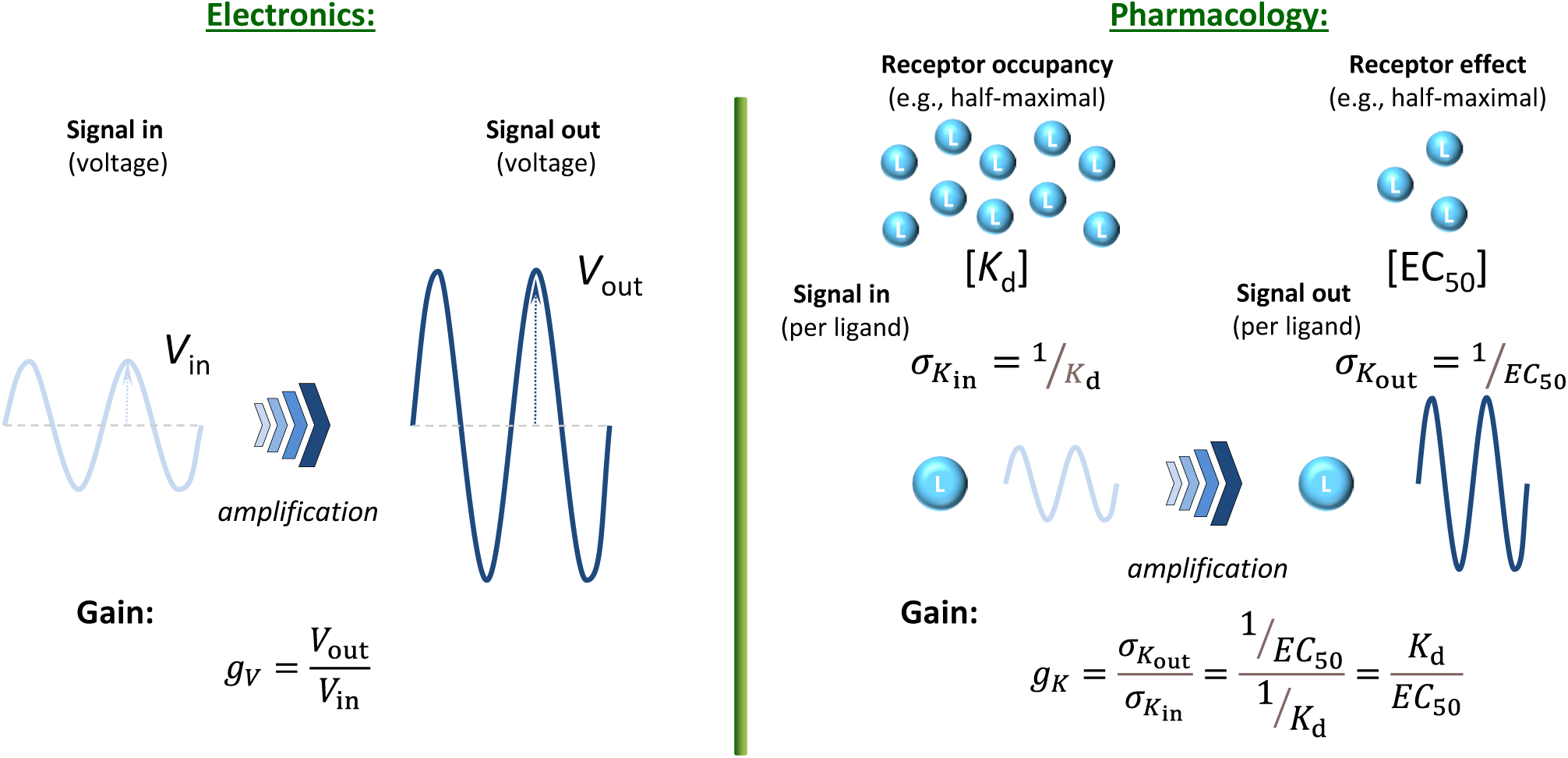
Quantification of signal amplification (gain) in pharmacology using *σ_K_* = 1/*K* (potency per ligand, i.e., effectiveness to generate a response) as signal. This way, a gain parameter can be defined using the half-maximal concentration values for occupancy (*K*_d_) and response (EC_50_) obtained with a full agonist, *g_K_* = *σ_K_*_out_/*σ_K_*_in_ = *K*_d_/EC_50_, and analogy with the gain parameter commonly used in electronics maintained.

### Gain Quantification Based on the Shift in Transfer Functions

One can also arrive at the same by considering that, as already mentioned, the classic Clark equation (or its more general form the Hill equation) serves well as transfer function (*f*_tr_, eq. 3) for most pharmacological responses (Figure 2). Thus, for normalized responses, the transfer function can be written as:

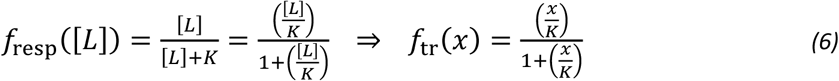

Rearranging eqs. 1 and 2 to have a similar form, they can be written as:

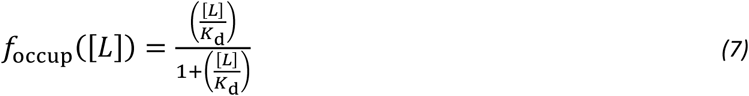

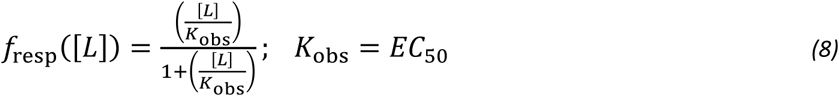

Comparing these, it is clear that the measured response, the “out” signal (eq. 8), follows the same pattern as the occupancy, the “in” signal (eq. 7), but with an amplified input *x* (assuming that the response runs “ahead” of occupancy, *K*_obs_ < *K*_d_). The ratio of these two *x*s can be considered as the gain factor of the corresponding signal transduction giving the same result as eq. 5:

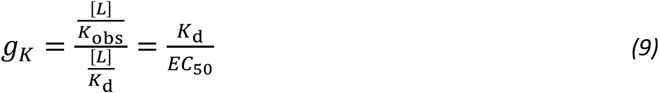

Not only is this definition of gain for receptor signal transduction (eq. 5 or 9) similar to that used in other fields such as electronics, but it also has the advantage that on the semi-log scales customarily used to plot concentration-response curves, it corresponds to the horizontal shift between log *K*_d_ and log *K*_obs_ (log EC_50_), so that it has an intuitive visual interpretation as well (Figure 4):

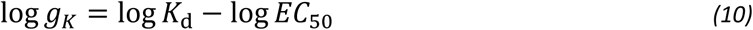

**Figure 4.**
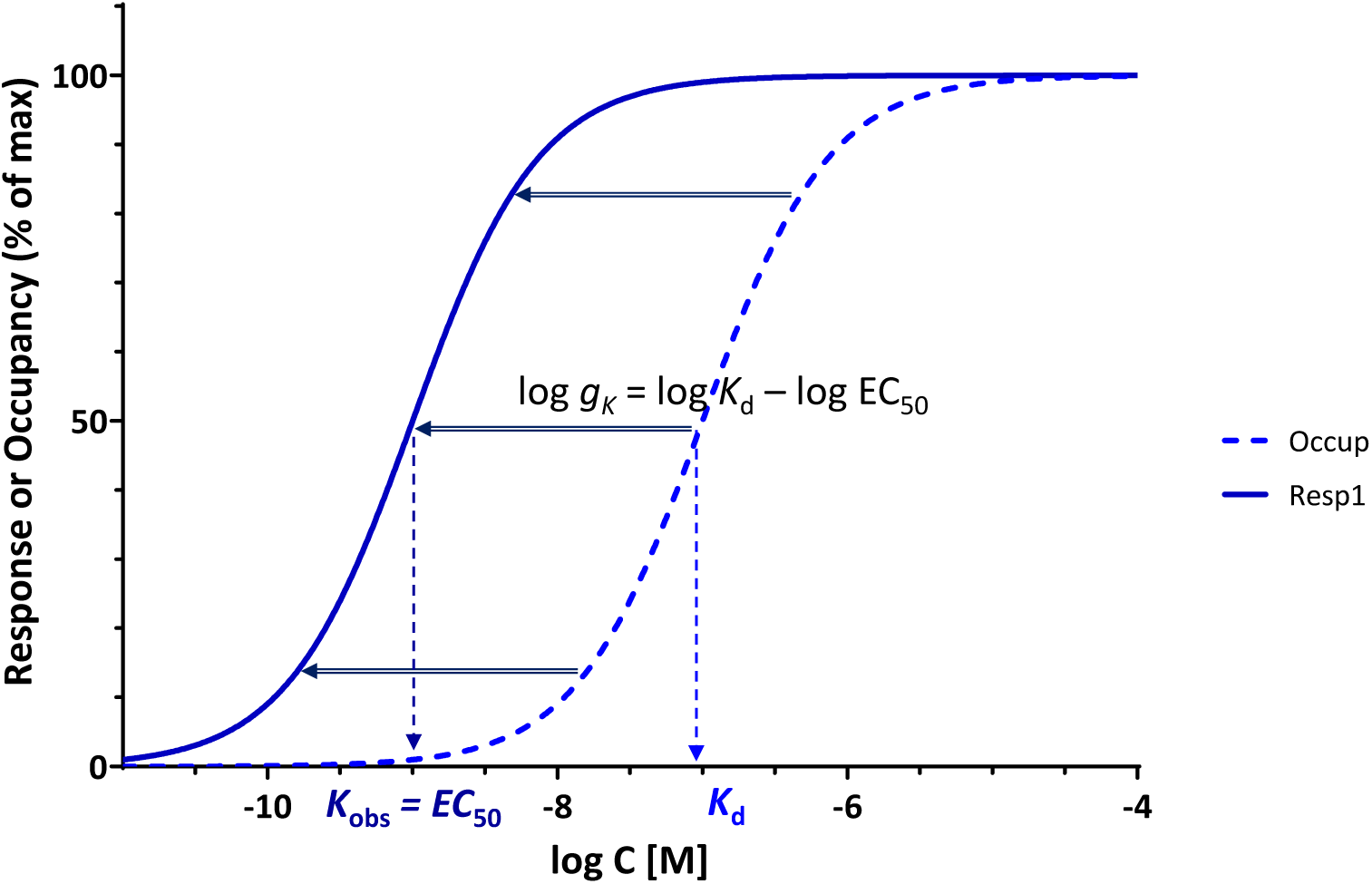
Use of *g_K_* = *K*_d_/EC_50_ of a full agonist as a parameter for gain quantification provides a convenient measure as it corresponds to the horizontal shift between the occupancy and response curves on typical semi-log graphs, log *g_K_* = log *K*_d_ – log EC_50_, a shift that for a full agonist is constant regardless of the point of assessment.

Furthermore, if both concentration-dependencies follow the same hyperbolic form (sigmoid on semi-log scale), a central assumption here, the horizontal shift between occupancy and response is the same everywhere as it is at the midpoint (*K*_d_ and EC_50_) as evident from comparing the functional forms in eq. 7 and 8 (see also detailed derivation for the general case in Supplementary Information, Appendix 1). Thus, log *g_K_*, as defined here is a particularly useful concept since for full agonists, the shift along the horizontal axis on the typical semi-log graph is the same everywhere, i.e., log *gκ* is the separation not just at the midpoint (at *K*_d_ and EC_50_ where *f*_occup_ = *f*_resp_ = 50%) but at any other arbitrary crosscut (e.g., at *f*_resp_ = *f*_occup_ = 25% or 90%; see Appendix 1, Supplementary Information). Note, however, that all these require full agonism, as response and occupancy need to run in parallel and both must reach 100% as maximum (Figure 4); partial agonists are discussed in the next section. Regarding this definition of gain for the presence of signal amplification (eq. 5), it should also be noted that it corresponds to what is generally considered as evidence for the existence of “*spare receptors*” (i.e., EC_50_ < *K*_d_) – see discussion later.

### Partial Agonist

Partial agonists cannot produce maximal response even at full occupancy. According to the official IUPHAR definition, a partial agonist is an “agonist that in a given tissue, under specified conditions, cannot elicit as large an effect (even when applied at high concentration, so that all the receptors should be occupied) as can another agonist acting through the same receptors in the same tissue” [31]. Thus, a partial agonist can only produce a maximum response (denoted as *e*_max_ on a normalized scale) that is less than that of the full agonist, *e*_max_ < 100%, even if occupancy has the same plateau of 100% receptor sites (denoted as *e*_100_). Hence, to accommodate partial agonists, eq. 2 needs to be modified to allow maximum responses that are less than 100% [8]:

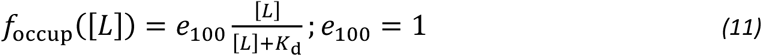

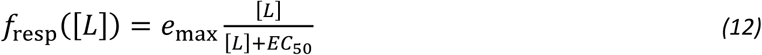

Note that for partial agonists, the half-maximal concentration *K*_obs_ is commonly denoted as maximum is produced (i.e., *e*_max_/2, which is less than 50%). Thus, if the partial agonist can only produce an *e*_max_ = 60% (compared to 100% for the full agonist), then its EC_50_ is at the concentration where it produces 30% response (*e*_max_/2) and not where it produces 50% response (see, e.g., Figure 5B for an illustration). The ratio of *K*_d_ and *K*_obs_ = EC_50_ will be denoted as *κ* for the general case (as it has been done before [8]):

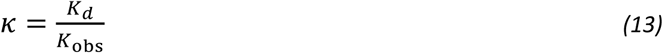

**Figure 5.**
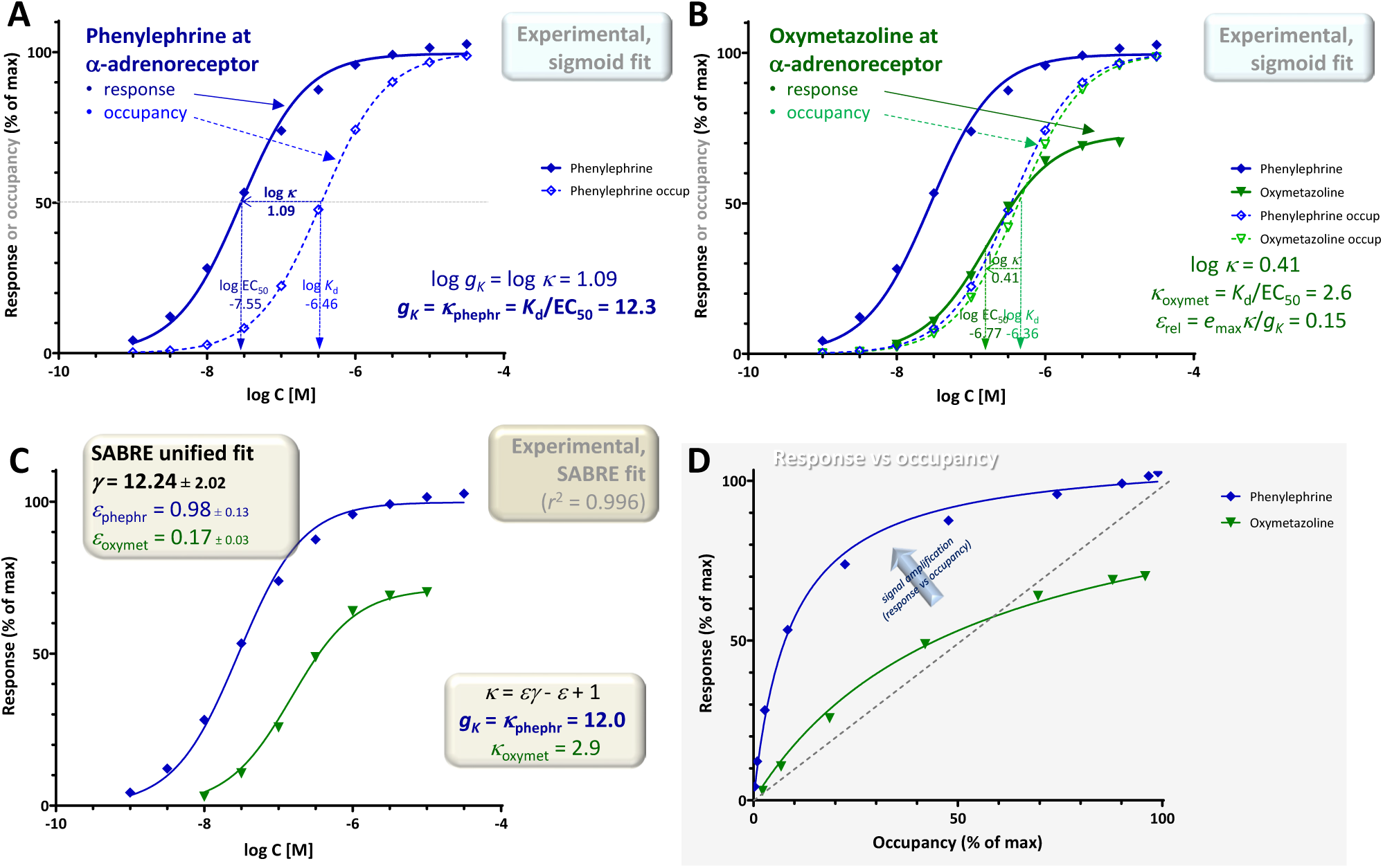
Concentration dependency of receptor occupancy (open symbols and dashed lines) and response (closed symbols and continuous lines) for phenylephrine (**A**, blue) and oxymetazoline (**B**, green), a full and a partial α-adrenoceptor agonist, respectively for experimental data (contractions of isolated rat aorta) [37] that is often used as textbook illustration (e.g., [6]). Corresponding *K*_d_ and EC_50_ estimates from fitting with classic hyperbolic equations (sigmoid on log-scale; eqs. 11-12) were used to obtain the *g_K_* = *κ*_phephr_ gain parameter (eq. 5) (**A**) as well as the *κ*_oxymet_ = *K*_d_/EC_50_ ratio (eq. 13) and *ε*_rel_ relative efficacy (eq. 16) for oxymetazoline (**B**). The same data was also fitted with SABRE (eq. 19) in a unified manner to determine the corresponding gain and efficacy parameters *γ*, *ε*_phephr_, and *ε*_oxymet_ (**C**). A graph of the response versus occupancy data is also included, fitted directly with the corresponding hyperbolic relationship between *f*_resp_ vs *f*_occup_ (eq. 26) (**D**). The gain parameter for this pathway obtained from the full agonist (phenylephrine) data alone *g_κ_* = 12.3 (**A**), is in excellent agreement with the global gain parameter *γ* = 12.24 ± 2.02 obtained from the fit of both the phenylephrine and oxymetazoline data by SABRE (**C**), which also suggests an efficacy *ε* = 0.17 ± 0.03 for oxymetazoline.

For partial agonists, *κ* will in general be smaller than that of the full agonist, which corresponds to the gain parameter for this response, *κ* < *κ*_full agon_ = *g_K_*, and it can be used to obtain an estimate of the relative efficacy. As shown before [17], for typical hyperbolic responses (sigmoid on semi-log scale), relative ligand efficacies can be compared using the *E*_max_·*K*_d_/EC_50_ ratios:

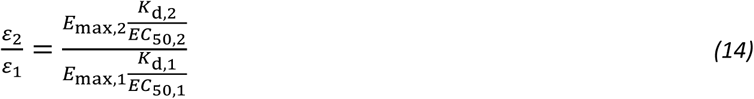

This formula was derived earlier for sigmoid responses from the assumption that at conditions that produce equal responses at low enough concentrations, the ratio of efficacies is the reverse of the ratio of occupied receptors producing it; see [17] for details. Use of the same formula to compare relative efficacies (*E*_max_·*K*_d_/EC_50_, eq. 14) has been also suggested by others based on different considerations [32-34]. Introducing *κ* and assuming normalized responses (with *e*_max_ maxima), eq. 14 with the present notation becomes:

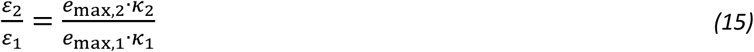

Thus, relative efficacies can be estimated by comparing *e*_max_·*κ* products. Consequently, for a partial agonist, its relative efficacy compared to the full agonist that produces 100% maximum response (and has an efficacy of one) can be obtained as:

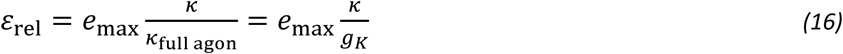

Finally, in addition to looking at responses as a function of ligand concentration, i.e., concentration-response curves, *f*_resp_ = ℱ([L]), it is also informative to look at them as a function of occupancy as well, *f*_resp_ = ℱ(*f*_occup_), especially in the present context of connecting receptor occupancy and response. Accordingly, for all examples below, a response versus occupancy graph will also be included for illustration. It has been shown before that if both occupancy and response follow hyperbolic relationships, as assumed here, fractional response (*f*_resp_) is a hyperbolic function of occupancy (*f*_occup_) even for partial agonists that can only produce a maximum response of *e*_max_ < 100% [8]:

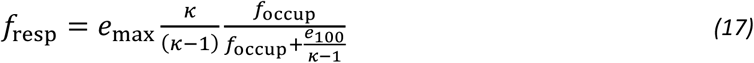

### Gain Quantification Using the SABRE Receptor Model

The gain quantification method discussed so far relies entirely on parameters derived from fitting the experimental data with standard concentration-response curves (*K*_d_, EC_50_) and the gain parameter *gκ* derived from them (eq. 5). Thus, it only relies on the assumption that both response and occupancy follow classic hyperbolic concentration dependencies (sigmoid on the commonly used semi-log scale) characterizable by the corresponding half-maximal concentrations values, *K*_d_ and EC_50_, respectively (eqs.11-12) and that they were determined for a true full agonist. Gain quantification of pharmacological signaling can also be done in a different, model-based approach using the recently introduced SABRE model (Signal Amplification, Binding affinity, and Receptor-activation Efficacy) – the first quantitative receptor model that explicitly includes parametrization for signal amplification [8, 18, 35]. SABRE, in its full general form, employs a total of five parameters to cover the full spectrum of parameters listed earlier in the Introduction – *K*_d_ for binding affinity, *ε* for efficacy, *ε*_R0_ for the efficacy of constitutive activity, *γ* for the gain of signal amplification, and *n* for Hill coefficient (for details, see [8, 18, 35]):

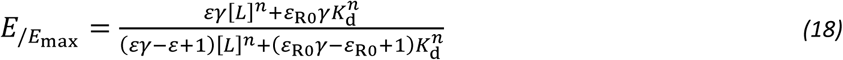

Here, only the simplified three-parameter version of SABRE will be used, which assumes that there is no constitutive activity (*ε*_R0_ = 0) and the regular law of mass action holds (Hill slope *n* = 1), but allows partial agonism (*ε*) and signal amplification (*γ*):

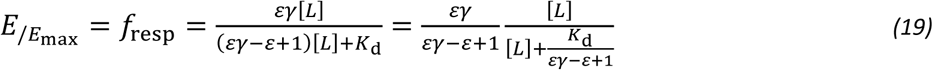

Comparing this equation (eq. 19) in its the rearranged form on the right side with eq. 12 describing the typical concentration-response curve, makes it clear that the assumptions of SABRE also result in a classic hyperbolic (sigmoid on the semi-log scale) relationship between (fractional) response, *E*_/*E*max_, and ligand concentration, [L], with apparent EC_50_ (*K*_obs_) and *e*_max_ (0 < *e*_max_ ≤ 100%) values that are:

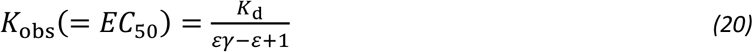

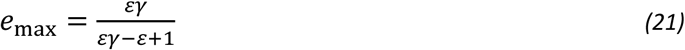

Accordingly, the *κ* ratio of *K*_d_/*K*_obs_ for a given agonist (eq. 13), can be expressed in terms of SABRE parameters as:

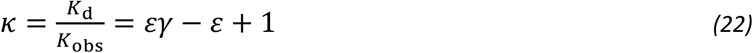

Notably, considering that for a full agonist (*ε* = 1), this *κ* ratio equals the gain parameter, *g_K_* = *κ*, it becomes obvious that *g_K_* corresponds exactly to the gain parameter *γ* of SABRE:

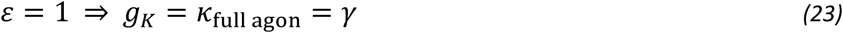

This is particularly encouraging, as it shows that the gain parameter (*g_K_*) obtained from concentration-response based considerations that led to eq. 5 and 9 corresponds to the gain parameter of SABRE (*γ*), which, however, was introduced based on different considerations (i.e., *γ* = [R_tot_]/*Kγ* to extend the range of the input of the hyperbolic response function linking the concentration of active receptors to response; see [18]). Accordingly, if data obtained with multiple agonists can be fitted adequately within the unified framework of SABRE (i.e., response data are adequately fitted while using the experimental *K*_d_ values as affinity parameters in SABRE, eq. 19), then the *γ* parameter of SABRE can provide a more reliable gain estimate than the *g_K_* value obtained from the full agonist data alone – several illustrative examples are provided below.

Furthermore, fit with SABRE also provides direct estimates of ligand efficacies, *ε*, for all agonists with response data included. In fact, if its assumptions hold, the relative efficacy of partial agonists derived earlier for hyperbolic responses (eq. 16) yields exactly the *ε* of SABRE using substitutions from eqs. 21-23:

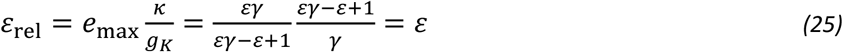

On the other hand, eq. 22 also provides a different formula to estimate *ε*_rel_ as long as the assumptions of SABRE are valid. Expressing *ε* as *ε*_rel_ of a partial agonist from eq. 22 (and using eq. 23 to replace *γ* with *g_K_*):

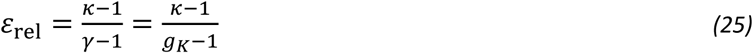

Finally, within the formalism of SABRE, the response versus occupancy relationship of eq. 17 can be written as:

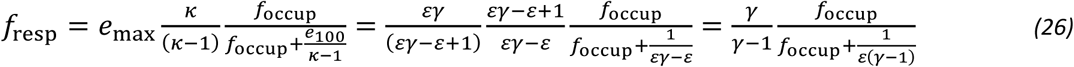

In the examples included in Results, this was used to fit the corresponding response versus occupancy curves.

### Data and Model Fitting

Experimental data and corresponding *K*_d_ and EC_50_ values used here are from published works as referenced for each case; response data values used for model fittings were obtained from the figures using WebPlotDigitizer [36]. All data used here were normalized to be in the 0–100% range and fitted using GraphPad Prism (GraphPad, La Jolla, CA, USA, RRID:SCR_002798). Fittings with SABRE were done with a custom implementation corresponding to the general eq. 19 (available for download, see [35]), and with parameters constrained as indicated for each case. Response versus occupancy data were fitted with a custom GraphPad model corresponding to eq. 26.

## Results and Discussion

A number of illustrative examples from published data involving both full and partial agonists acting on various receptors are provided to illustrate the gain quantification process described above. There is only relatively limited data where both receptor binding (occupancy) and response were measured in parallel in the same system as needed here; nevertheless, the examples below should be sufficient to support the concept of signal amplification at pharmacological receptors and the advantages of using *g_K_* = *K*_d_/EC_50_ as a gain parameter for its quantification.

### Example 1: Single Receptor (α-Adrenergic), Pathway, and Readout (Rat Aorta Contraction) with Multiple Agonists

A first illustration is provided with data that is frequently used as textbook illustration of the possible complex relationship between receptor occupancy and response (for example, in *Rang and Dale’s Pharmacology* [6]): the concentration-dependent contractions of isolated rat aorta induced by imidazoline-type α-adrenoceptor agonists including phenylephrine, oxymetazoline, and others [37]. Contractions of isolated rat aortic strips were measured as response using isometric transducers, and receptor binding affinities (dissociation constants, *K*_d_s) were assessed separately by two different methods using Furchgott-type [16] and Schild-plot based methods [38], respectively [37]; the average of these two affinity estimates were used here. From the whole dataset obtained originally with ten compounds [37], here, only the data of phenylephrine as full agonist and oxymetazoline as representative partial agonist are used to avoid overcrowding the figures (Figure 5). Experimental data obtained with the full agonist phenylephrine (log *K*_d_ = –6.46, log EC_50_ = –7.55), indicate a shift between the semi-log curves of response and occupancy of log *κ* = log *K*_d_ – log EC_50_ = 1.09; thus a corresponding gain parameter for this readout that is *g_K_* = *κ*_full agon_ = *K*_d_/EC_50_ = 12.3 (Figure 5A). The shift between the response and occupancy curves for the partial agonist is smaller; here, log *κ* = 0.41 (green in Figure 5B with log *K*_d_ = –6.36, log EC_50_ = –6.77), thus *κ* = *K*_d_/EC_50_ = 2.6 for oxymetazoline, as the response it generates is running less “ahead” of occupancy than for the full agonist (is less left-shifted) as clearly noticeable in Figure 5D. In fact, oxymetazoline illustrates nicely one of the confusing cases when comparing (fractional) response and occupancy for a partial agonist acting on a pathway with amplification as its response actually runs both “ahead” and “behind” the occupancy: *f*_resp_ > *f*_occup_ until about 60% occupancy (*f*_occup_ < 0.6), but then reverses and *f*_resp_ < *f*_occup_ when *f*_occup_ > 0.6 (Figure 5D). According to this data, using eq. 16, the relative efficacy of oxymetazoline compared to phenylephrine is *ε*_rel_ = 0.73×2.6/12.3 = 0.15 (Figure 5B).

These estimates (signal transduction gain *g_K_* = 12.3 and relative efficacy of oxymetazoline *ε*_rel_ = 0.15) are solely based on fitting of the experimental data with individual sigmoid response curves and the corresponding *K*_d_ and EC_50_ values. As the data in its entirety can be fitted with SABRE as a single, unified model (see [18] for a detailed fit including for multiple partial agonists), estimates can also be obtained from fit of the entire data with a single set of model parameters. Overall, SABRE (eq. 19) fits the data very well accounting for more than 99% of the variability (*r*^2^ = 0.996; Figure 5C). Unified fitting of the entire dataset results in a gain parameter, *γ* = 12.24 ± 2.02, and an efficacy for oxymetazoline, *ε* = 0.17 ± 0.03, that are both in excellent agreement with the estimates obtained simply from the experimental *K*_d_ and EC_50_ values using eq. 5 and 16, respectively indicating that very consistent estimates can be obtained for this data.

### Example 2: Single Receptor (M_3_ Muscarinic) and Pathway with Multiple Readouts (G_α_-GTP Binding & Intracellular Ca Increase)

A second illustration is provided with a dataset involving responses by the M_3_ muscarinic receptor elicited by agonists including oxotremorine-M and methacholine [39]. It is included as an illustration for a case where two different responses are measured at consecutive vantage points downstream on the same pathway: here, the stimulation of GTP binding to G_α_ and the subsequent increase in intracellular calcium, Ca (Figure 6; shown as lighter half-closed and darker closed symbols, respectively). GTP binding assays were performed using [^35^S]GTPγS in 96-well optiplates, agonist-induced changes in Ca^2+^ concentration were measured using a fluorometric imaging plate reader, and binding affinity estimates (*K*) were obtained from equilibrium competition experiments with *N*-methyl-[^3^H]scopolamine (following Cheng-Prusoff corrections [40]) [39]. As immediately evident from a quick comparison of the log *K*_d_ and log EC_50_ values in Figure 6A, the amplifications of the two responses assessed here are very different with that of the GTP binding being close to unity (log *κ*_GTP_ = 0.20; *g_K_*_,GTP_ =*κ*_GTP,full agon_ = 1.6) and that of the Ca increase being very large, around four orders of magnitude (log *κ*_Ca_ = 4.15; *g_K_*_,Ca_ = *κ*_Ca,full agon_ ≈ 14000). Such a large discrepancy makes it difficult to obtain well-defined fittings and parameter estimates, but the partial agonist character of methacholine is clearly observable in the GTP response (*e*_max_ = 0.66) and the decreased shift in the Ca response (*ε*_rel,Ca_ = 0.71) (Figure 6B).

**Figure 6.**
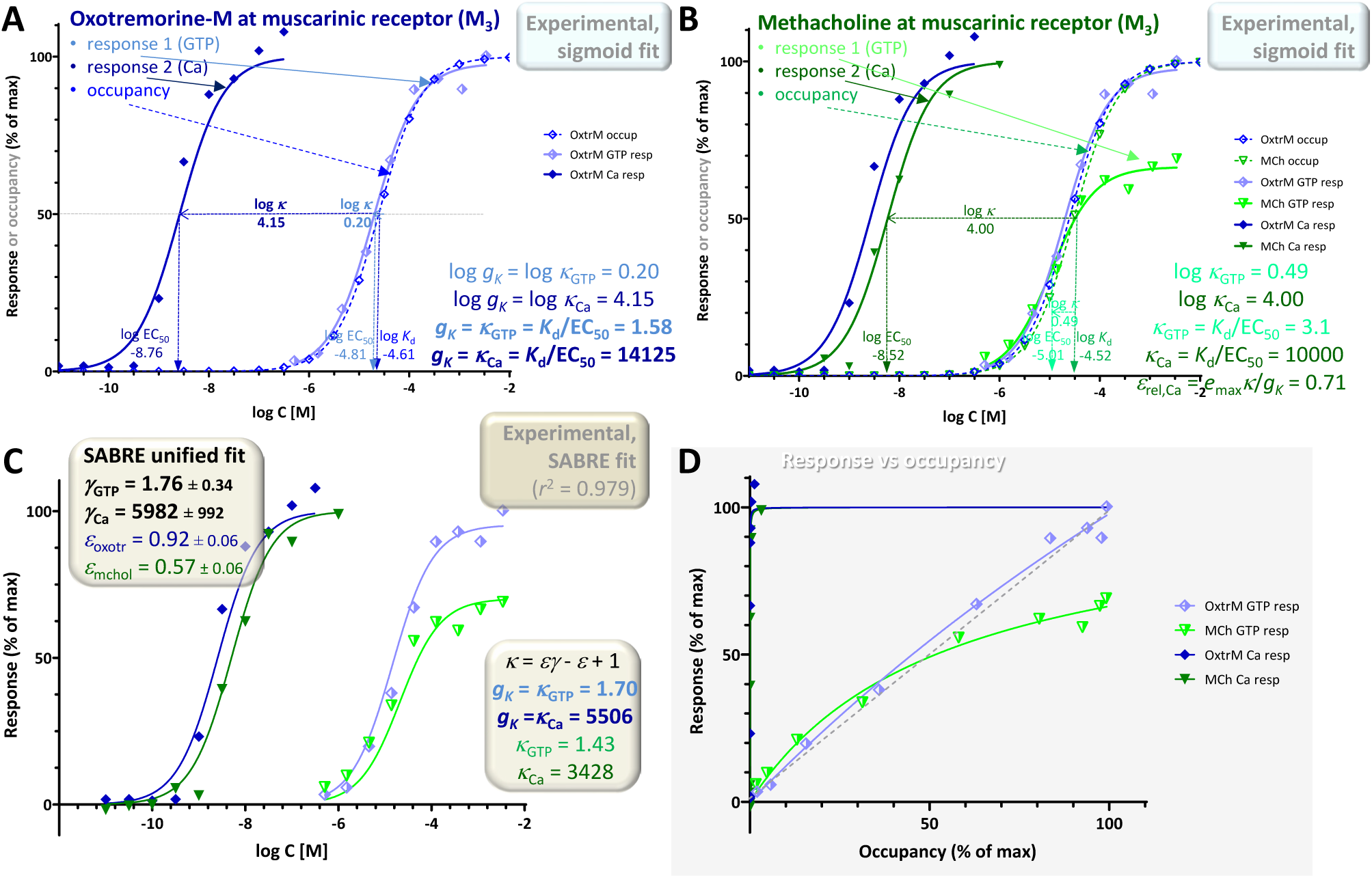
Concentration dependency of receptor occupancy (open symbols and dashed lines) and two different responses (stimulation of GTP binding to Gα subunits and subsequent increase in intracellular Ca levels after M_3_ receptor activation; lighter half-closed and darker closed symbols, respectively with continuous lines) for the muscarinic agonists oxotremorine-M (**A**, blue) and methacholine (**B**, green). Experimental data [39] and the corresponding *K*_d_ and EC_50_ estimates from fitting with classic hyperbolic equations (sigmoid on log-scale; eqs. 11-12) were used to obtain the two different *g_K_* gain parameter (eq. 5) (**A**) and *κ* values (eq. 13) (**B**). Fit of the full agonist (oxotremorine-M) data alone indicates gains *g_K_* of 1.6 and ∼14000 for GTP and Ca, respectively (**A**). As in Figure 5, the data was also fitted with SABRE (eq. 19) in a unified manner to determine the corresponding gain (*γ*_GTP_, *γ*_Ca_) and efficacy parameters (*ε*_oxotr_, *ε*_mchol_) (**C**). Values obtained this way for the gain parameters (**C**) are in general agreement with those from the sigmoid fit (**A**); they also suggest an efficacy *ε* = 0.57 ± 0.06 for methacholine. A graph of the response versus occupancy data is also included, fitted directly with the corresponding hyperbolic relationship between *f*_resp_ vs *f*_occup_ (eq. 26) (**D**).

For the same reason, fit of the entire dataset with SABRE (eq. 19) as a single model is particularly challenging; nevertheless, it still accounts for most of the variability (*r*^2^ = 0.979; Figure 6C) suggesting gain (amplification) parameters (*γ*_GTP_ = 1.76 ± 0.34, *γ*_Ca_ = 5982 ± 992) and an efficacy for the partial agonist methacholine (*ε* = 0.57 ± 0.06) in reasonable agreement with those from the previous sigmoid-based estimates. Due to the strong amplification in the Ca responses, the corresponding response versus occupancy curve is strongly distorted as even minimal occupancy already results in essentially maximum response (Figure 6D). Nevertheless, these data still provide an illustration of a case with two different responses measured along the same downstream pathway with two different amplifications corresponding to the two different assessment points. It also highlights an issue that often causes confusion: partial agonists are typically recognized as not being able to elicit maximum responses; however, in pathways with strong amplification, even relatively weak partial agonists can cause full responses as nicely evidenced here by the Ca response of methacholine (versus the corresponding GTP response; closed vs half-closed green symbols in Figure 6). A more detailed illustration of the effect of intermixing the effect of partial agonism and different amplification / receptor levels is shown in Figure 7 and discussed below.

**Figure 7.**
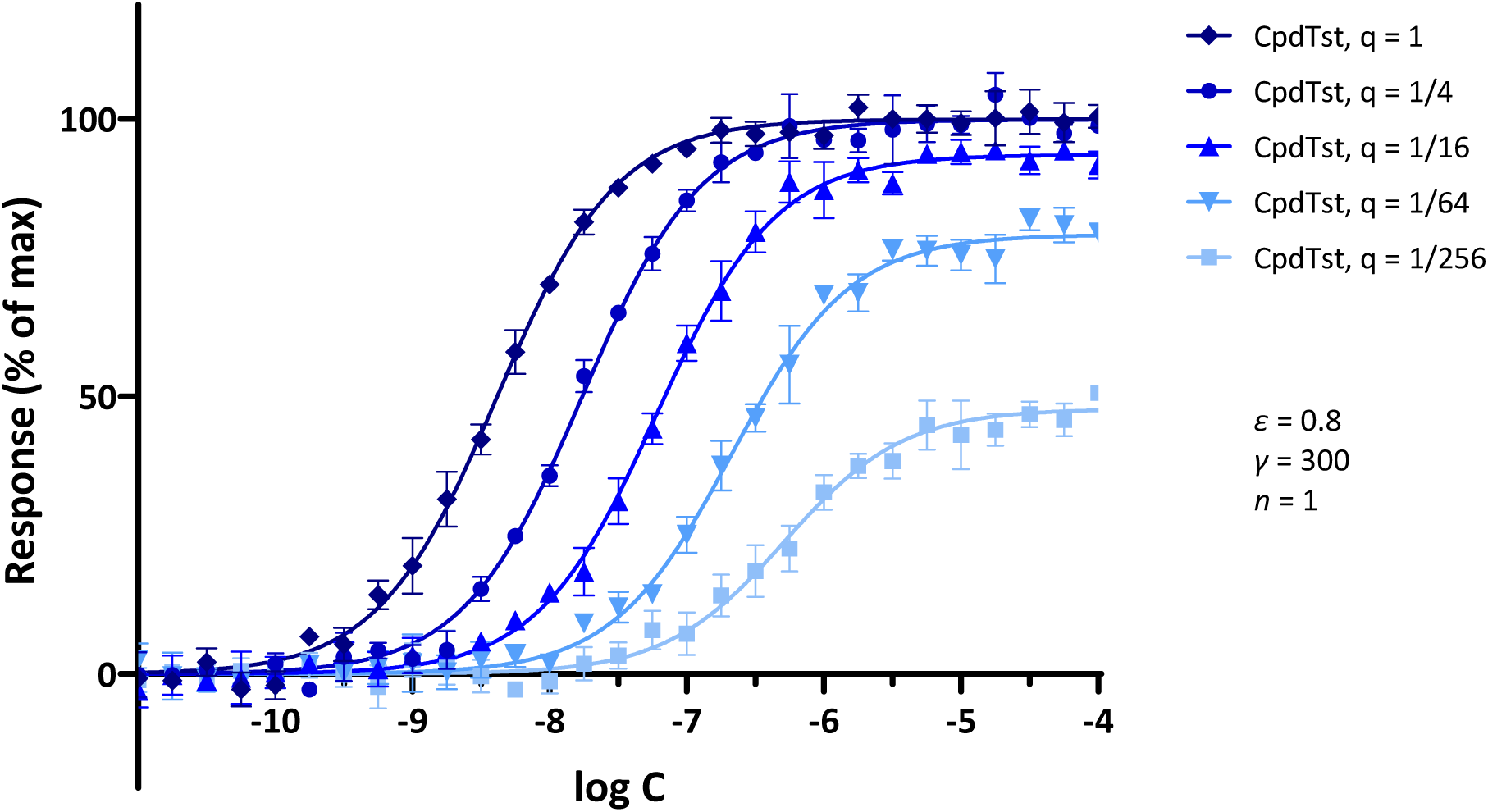
Illustrative concentration-response curves generated for a hypothetical agonist following increasing levels of partial irreversible receptor inactivation (Furchgott method). Simulated data were generated with SABRE for a hypothetical ligand (CpdTst; *K*_d_ = 1 μM, *ε* = 0.8) and pathway gain (*γ* = 300; 5% random error) assuming consecutive four-fold irreversible inactivations (as indicated by the *q* values of 1, 1/4, 1/16, 1/64, and 1/256 for the remaining fractions of receptors; see [17] for model details). Data were selected so as to reproduce a textbook illustration of the method used to demonstrate experimentally the presence of *spare receptors* (see [7]) and to show that such response-profiles can also be obtained assuming signal amplification and consecutive reduction in receptor numbers. Model parameters were also selected to illustrate that if there is sufficient amplification, full maximal response can be achieved even with a partial agonist (*ε* = 0.8) and even after inactivation of a considerable fraction of receptors (e.g., *q* = 1/4); however, if sufficient receptors are inactivated, maximal responses will become diminished.

### Example 3: Single Receptor (Muscarinic), Pathway, and Readout (Adenylate Cyclase, Rabbit Myocardium) with Multiple Levels of Partial Irreversible Inactivation (Furchgott Method)

The third illustration is for a set of Furchgott type experiments that allow the quantification of receptor binding affinity by comparing concentration-response curves obtained at different receptor levels, e.g., following partial irreversible inactivation. One advantage of such experiments is that they require the measurement of response data only and use that to estimate binding affinity; thus, they avoid the need of having to set up separate ligand binding experiments [16, 17]. A set of illustrative concentration-response curves obtained from simulations of such an experiment assuming increasing levels of partial irreversible receptor inactivation is shown in Figure 7. Simulation parameters were selected so as to reproduce textbook illustrations of the Furchgott method used to demonstrate experimentally the presence of *spare receptors* (e.g., [7]). The data from such multiple concentration-response curves can be used to estimate both the *K*_d_ of the ligand and the fraction of remaining receptors after each inactivation (*q*) [17]. Thus, such Furchgott-type approaches are particularly well-suited for quantitative pharmacology characterization as they allow the concurrent estimation of the binding affinity and relative efficacy of the ligand as well as the characteristics of the pathway as long as multiple concentration-response curves can be obtained at different levels of receptor levels [17] either via the classic method of partial irreversible receptor inactivation as introduced by Furchgott [15, 16] or via different levels of receptor expression, which are now possible (see, e.g., [41] for a recent implementation).

The experimental data used for illustration here were all also obtained using muscarinic receptors and oxotremorine-M as full agonist as in example 2 but involve muscarinic receptor-mediated inhibition of adenylate cyclase activity as response (Figure 8). They were obtained in perfused rabbit myocardium homogenates following different levels of partial inactivation with an irreversible muscarinic antagonist (benzilylcholine mustard, BCM) [42]. Responses obtained with the full agonist (oxotremorine-M) and no inactivation give a shift between the response and occupancy curves of log *κ* = 1.00 corresponding to a gain of *g_K_* = *κ*_full agon_ = *K*_d_/EC_50_ = 10.0 (log *K*_d_ = –5.40, log EC_50_ = –6.40; Figure 8A). Following partial inactivations, the shifts become smaller: 0.54 and 0.25 for the two different levels used here (BCM at 1 and 10 nM for 15 min [42]; Figure 8B). The fit of these curves with individual sigmoid curves suggest a log *K*_d_ of –5.35 for oxotremorine-M (in excellent agreement with the value measured by a competition assay, –5.40) as well as remaining fractions of receptors of *q*_1_ = 0.29 and *q*_2_ = 0.06, respectively (Figure 8B; see also Table S4A in [17]). The response versus occupancy graph (Figure 8D) nicely shows the effect of these inactivations and corresponding irreversible reductions in receptor levels on the fractional responses achievable at various occupancy levels.

**Figure 8.**
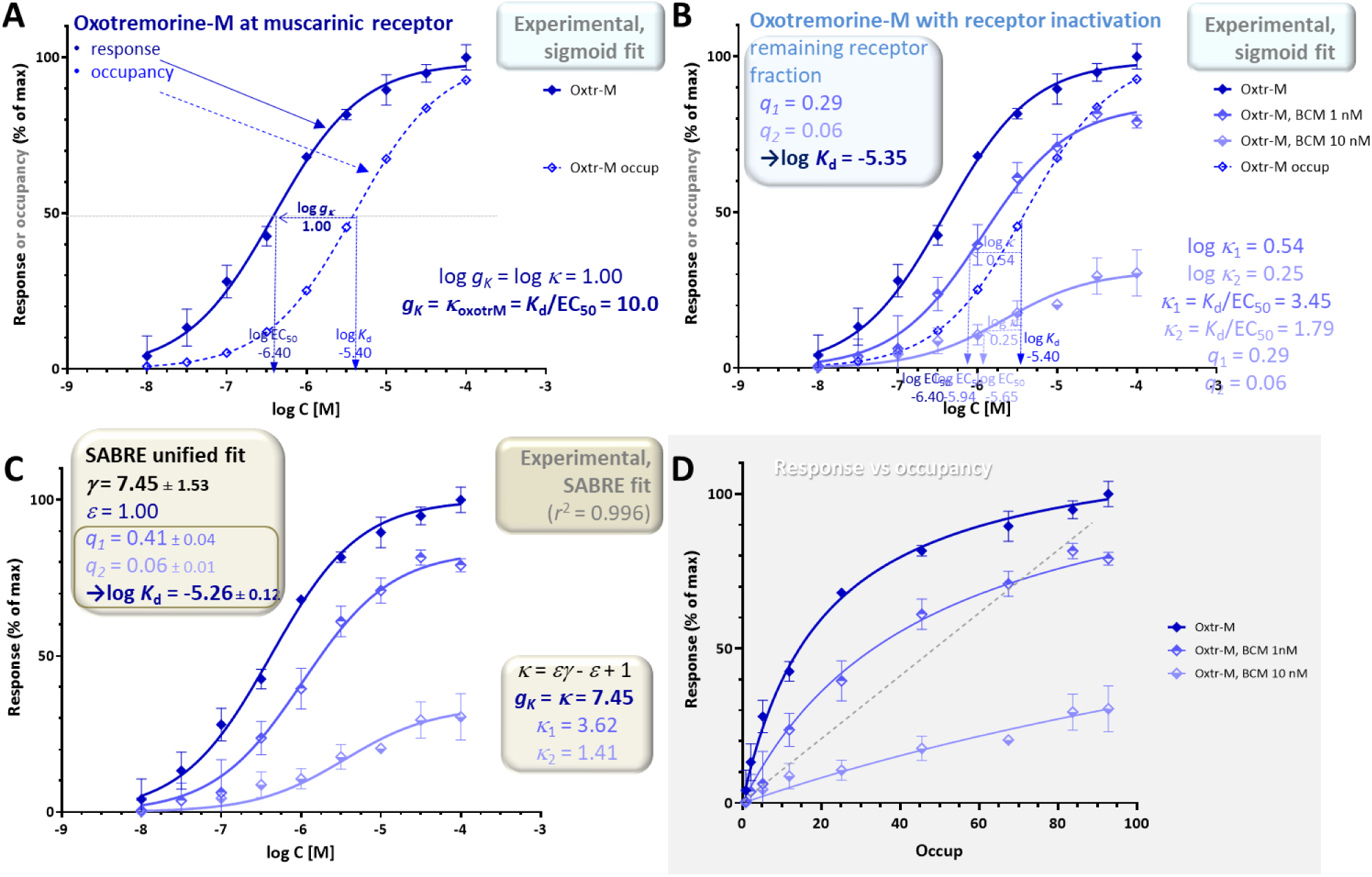
Concentration dependency of receptor occupancy (open symbols and dashed lines) and response (muscarinic receptor-mediated inhibition of adenylate cyclase activity in perfused rabbit myocardium; closed symbols and continuous line) for oxotremorine-M (**A**, dark blue) and the same following two different levels of partial irreversible inactivation (**B**, lighter blue colors). Experimental data [42] and the corresponding *K*_d_ and EC_50_ estimates were used to obtain the *g_K_* gain parameter (**A**) and *κ* values (**B**) as before. The entire dataset was also fitted with SABRE to determine the gain parameter and the remaining fraction of receptors resulting in *γ* = 7.45 ± 1.53, *q*_1_ = 0.41 ± 0.04, and *q*_2_ = 0.06 ± 0.01 (**C**). A graph of the response versus occupancy data is also included, fitted directly with the corresponding hyperbolic relationship between *f*_resp_ vs *f*_occup_ (eq. 26) (**D**).

As in the previous examples, the entire data can also be fitted with SABRE using a single set of parameters to obtain unified model-based estimates. For this data, SABRE gives excellent overall fit (*r*^2^ = 0.996; Figure 8C), and the unified fitting of all three curves result in a gain parameter of *γ* = 7.45 ± 1.53 for this response and log *K*_d_ = –5.26 ± 0.12 for oxotremorine-M – in good agreement with the previous sigmoid fit based estimates as well as the measured value of log *K*_d_. With SABRE, the inactivation-caused fold decreases in receptor level show up as apparent fold reduction in efficacy (*ε*’=*qε*; see [17] for details); the corresponding values obtained here are *q*_1_ = 0.41 ± 0.04 and *q*_2_ = 0.06 ± 0.01, which are again consistent with the sigmoid-based estimates.

### Example 4: Single Receptor (AT1R Angiotensin), Two Different Pathways and Readouts (G_q_-Mediated Inositol Monophosphate Increase and β-Arrestin2 Endocytosis) with Balanced and Biased Agonists

The fourth illustration involves two different pathways originating from the same receptor, the angiotensin II receptor 1 (AT1R), but mediated by a G-protein and β-arrestin, respectively. Signal amplification can be different along such divergent pathways, so that even if the activation signal is the same, responses can be different – this is sometimes designated as “system bias”. It is also possible that, even if these pathways originate from the same receptor, ligands can activate them differently – a phenomenon termed as biased agonism (functional selectivity) (Figure 9) [43-47]. For biased agonism to exist, there must be different active states of the receptor that can preferentially initiate either one or the other downstream signal, and there must be agonists that can differentially stabilize these active states. Biased agonism is of considerable interest as it might allow improved therapeutic action by separating the desired activity from the unwanted side effects that are mediated by separate pathways, which is not possible with classical full or partial agonists (Figure 9). Biased agonism is, however, difficult to quantify [48-50]. For example, it is possible that even oliceridine, which was developed as a μ-opioid receptor (MOPr) biased agonist [51] and is being often touted to illustrate the clinical promise of biased agonists since its approval by the FDA (Olinvyk, 2020), is not a biased agonist but a regular weak agonist acting on one amplified and one unamplified or possibly attenuated pathway [8, 52].

**Figure 9.**
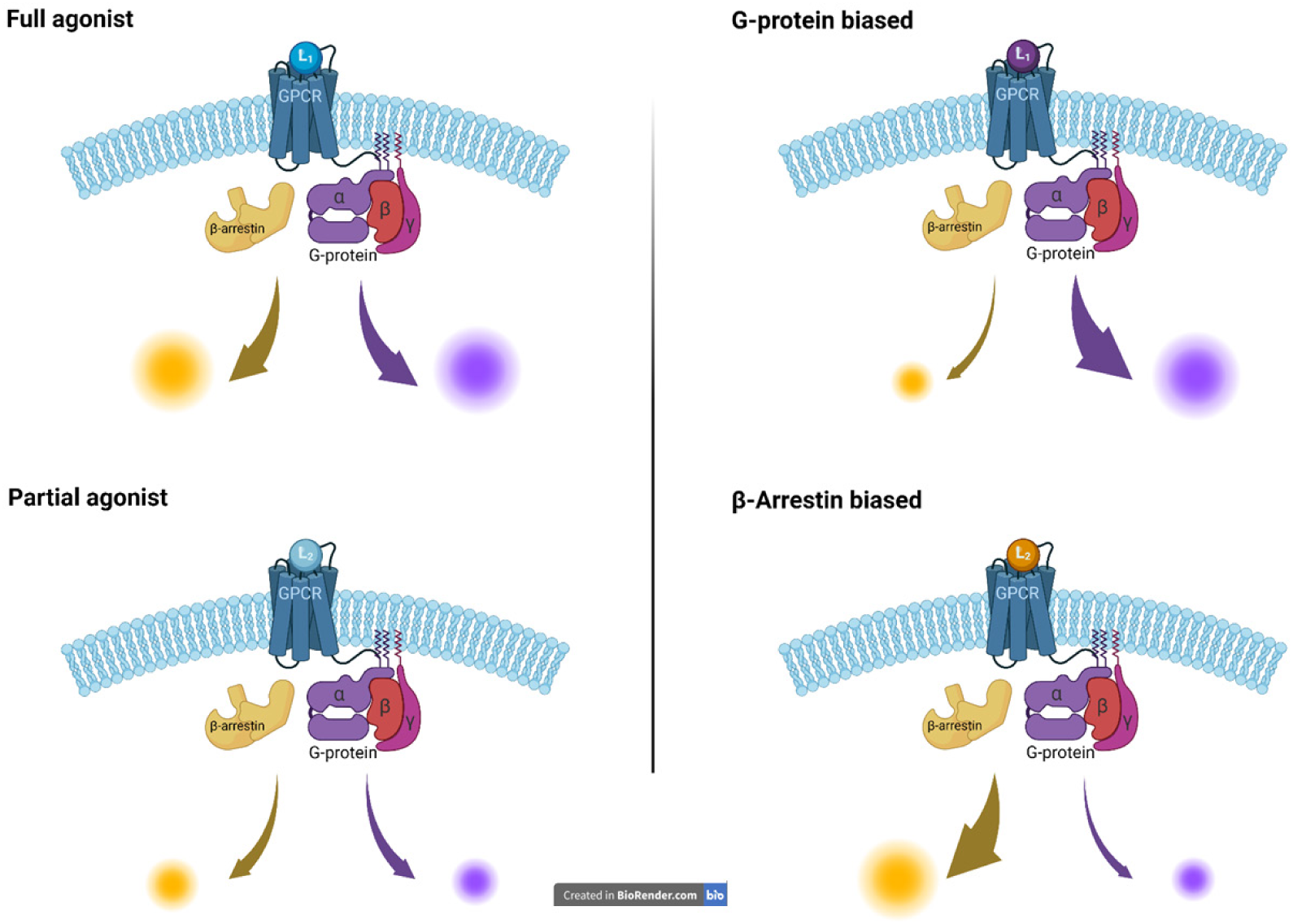
Illustration of the concept of biased agonism, which can achieve different activations along different pathways even if they originate from the same receptor (right column), compared to that of classic full and partial agonism, which activate all downstream pathway to the same full or partial degree (left column). A case of a G-protein coupled receptor (GPCR) is shown here with downstream signaling along a G-protein and a β-arrestin modulated pathway (figure created in BioRender).

Experimental data used for this example are for two responses, G_q_-mediated inositol monophosphate increase and β-arrestin2 endocytosis, generated by angiotensin II and TRV023 as agonists of AT1R (Figure 10) [53]. Responses were measured using the IP-One Gq kit from Cisbio and the PathHunter assay from DiscoverX, respectively, while binding affinities (*K*_d_ values) were determined in equilibrium competition radioligand binding assays with [^3^H]-olmesartan [53]. Fit of the sigmoid responses for the full agonist angiotensin II indicate gains *g_K_* = *κ_full agon_* of 10.2 and 6.6 for the G-protein and β-arrestin responses, respectively (Figure 10A). The same *κ* = *K*_d_/EC_50_ ratios for the partial agonist TRV023 are 0.25 and 7.1, respectively indicating a notably weaker G-protein response (Figure 10B). Accordingly, biased agonism for TRV023 is clearly evident in the response versus occupancy graph, where the G-protein response significantly deviates from the β-arrestin one for TRV023, whereas it does not for angiotensin II (Figure 10D). In agreement with this, the relative efficacies for TRV023 (eq. 16) for the two pathways, *ε*_Gprt_ = 0.0017 and *ε*_βArr_ = 0.22 indicate an approximately 13-fold difference.

**Figure 10.**
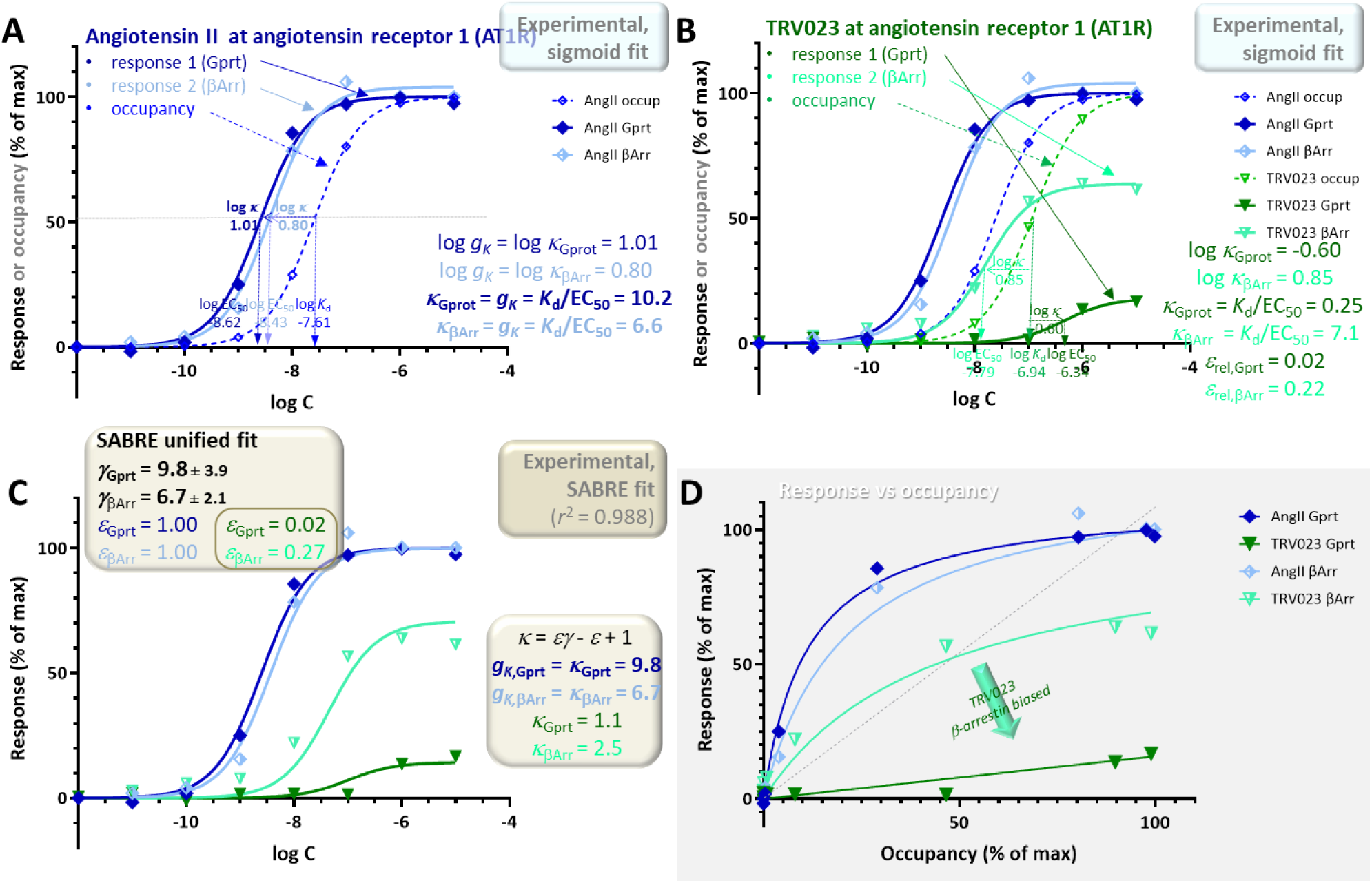
Concentration dependency of receptor occupancy (open symbols and dashed lines) and two different responses measured along different downstream pathways (G_q_-mediated inositol monophosphate increases and β-arrestin2 endocytosis – darker closed and lighter half-closed symbols, respectively with continuous lines) at the angiotensin II type 1 receptor (AT1R) for angiotensin II (**A**, blue) and TRV023 (**B**, green), a full and a partial agonist, respectively. Experimental data [53] and the corresponding *K*_d_ and EC_50_ estimates were used to obtain the two different *g_K_* gain parameters (eq. 5) (**A**) and *κ* values (eq. 13) (**B**). Sigmoid fit of the full agonist angiotensin II data indicates gains *g_K_* of 10.2 and 6.6 for the G-protein and β-arrestin responses, respectively (**A**). Estimates of the relative efficacies (eq. 16) for TRV023, *ε*_Gprt_ = 0.017 and *ε*_βArr_ = 0.22, indicate an about 13-fold difference suggesting β-arrestin biased agonism. As before, the same data was also fitted with SABRE (eq. 19) in a unified manner to determine the corresponding gain (*γ*_Gprot_, *γ*_βArr_) and efficacy parameters (*ε*_Gprot,AngII_, *ε*_βArr,AngII_, *ε*_Gprot,TRV_, *ε*_βArr,TRV_) (**C**). The gain parameters obtained from the fit of all data by SABRE (*γ*_Gprot_ = 9.8 ± 3.9 and *γ*_βArr_ = 6.7 ± 2.1) are in good agreement with those from the sigmoid fit (**A**). For TRV023, the SABRE calculated efficacies also clearly indicate β-arrestin biased agonism with an about 15-fold different in efficacies (*ε*_Gprt_ = 0.02 ± 0.008 vs *ε*_βArr_ = 0.27 ± 0.07), which is also evident in the response versus occupancy graph (**D**).

Such multi-pathway responses can also be fitted within the unified framework of SABRE using a single gain parameter *γ* _P_ for each pathway (P*_i_*) and a single experimental *K*_d_ but different pathway-dependent efficacies *ε*_P, L_ for each ligand (L*_j_*) [17, 18]. Such unified fit of the present data with this six-parameter model (*γ*_Gprt_, *γ*_βArr_, *ε*_Gprt,Ang_, *ε*_βArr,Ang_, *ε*_Gprt,TRV_, *ε*_βArr,TRV_) gives good fit (*r*^2^ = 0.988; Figure 10C), and the gain parameters obtained from the fit of all data with SABRE (*γ*_Gprt_ = 9.8 ± 3.9 and *γ*_βArr_ = 6.7 ± 2.1) are in general agreement with the *g_K_* pathway amplifications obtained from the *K*_d_/EC_50_ ratios of the full agonist angiotensin II (Figure 10). For TRV023, the SABRE calculated efficacies indicate clear β-arrestin biased agonism with an about 15-fold different in efficacies: *ε*_Gprt,TRV_ = 0.02 ± 0.008 vs *ε*_βArr,TRV_ = 0.27 ± 0.07.

### Example 5: Single Receptor (μ-Opioid), Two Different Pathways and Readouts (G_αi2_ Activation and β-Arrestin2 Recruitment) with Both Left- and Right-Shifted Responses

A fifth example included here also involves G-protein and β-arrestin mediated diverging pathways, but this time with responses initiated at the μ-opioid receptor (MOPr) by DAMGO (D-Ala^2^, is included to illustrate the unusual case where response curves are right- and not left-shifted compared to occupancy (meaning *κ* = *K*_d_/EC_50_ < 1). In the present context, this indicates not signal amplification, but apparent signal attenuation/dampening or loss (*g_K_* < 1) [8]. Experimental data were obtained in HEK293A cells using BRET assays to measure G_αi2_ activation as well as β-arrestin2 recruitment [54]. Binding measurements were done with [^3^H]naloxone, and equilibrium dissociation constants, *K*_d_, were calculated using the Cheng-Prusoff equation [40] to account for radioligand concentration [54]. Notably, while both responses follow classic hyperbolic shapes, and the G-protein mediated responses are left-shifted as typical for signal amplification cases as discussed here, the β-arrestin responses are clearly right-shifted compared to occupancy despite originating from the very same receptor (Figure 11). This is evident in the classic concentration-response curves (Figure 11A,B) as well as in the response versus occupancy graph, where they clearly have a different curvature (Figure 11D). The β-arrestin response is clearly lagging the occupancy even for the full agonist (DAMGO) while the G-protein response is ahead (dark vs light blue curves). For example, at 25% receptor occupancy, the G-protein response is already plateauing as it is approaching its maximum, whereas the β-arrestin response is minimal and still barely separated from baseline (*f*_occup_ = 25% → *f*_resp,Gprot_ > 80%, *f*_resp,βArr_ < 5% for DAMGO in Figure 11D). Thus, for these MOPr responses, EC_50,Gprt_ < *K*_d_ < EC_50,βArr_, so that the G-protein responses are left-shifted compared to occupancy indicating signal amplification (*g_K_*_,Gprot_ = *κ*_full agon_ = *K*_d_/EC_50,Gprt_ > 1), but the β-arrestin responses are right-shifted (less concentration sensitive), indicating the opposite (*g_K_*_,βArr_ = *κ*_full agon_ = *K*_d_/EC_50,βArr_ < 1). Indeed, sigmoid fit of the DAMGO data gave *g_K_*_,Gprot_ = 18.6 and *g_K_*_,βArr_ = 0.06 (Figure 11A). Notably, this unusual behavior of one left- and one right-shifted response is not just the case for the data shown here [54] as very similar results have been obtained in two other independent works with both DAMGO and morphine [55, 56] (see [8] for more detailed comparisons and discussions).

**Figure 11.**
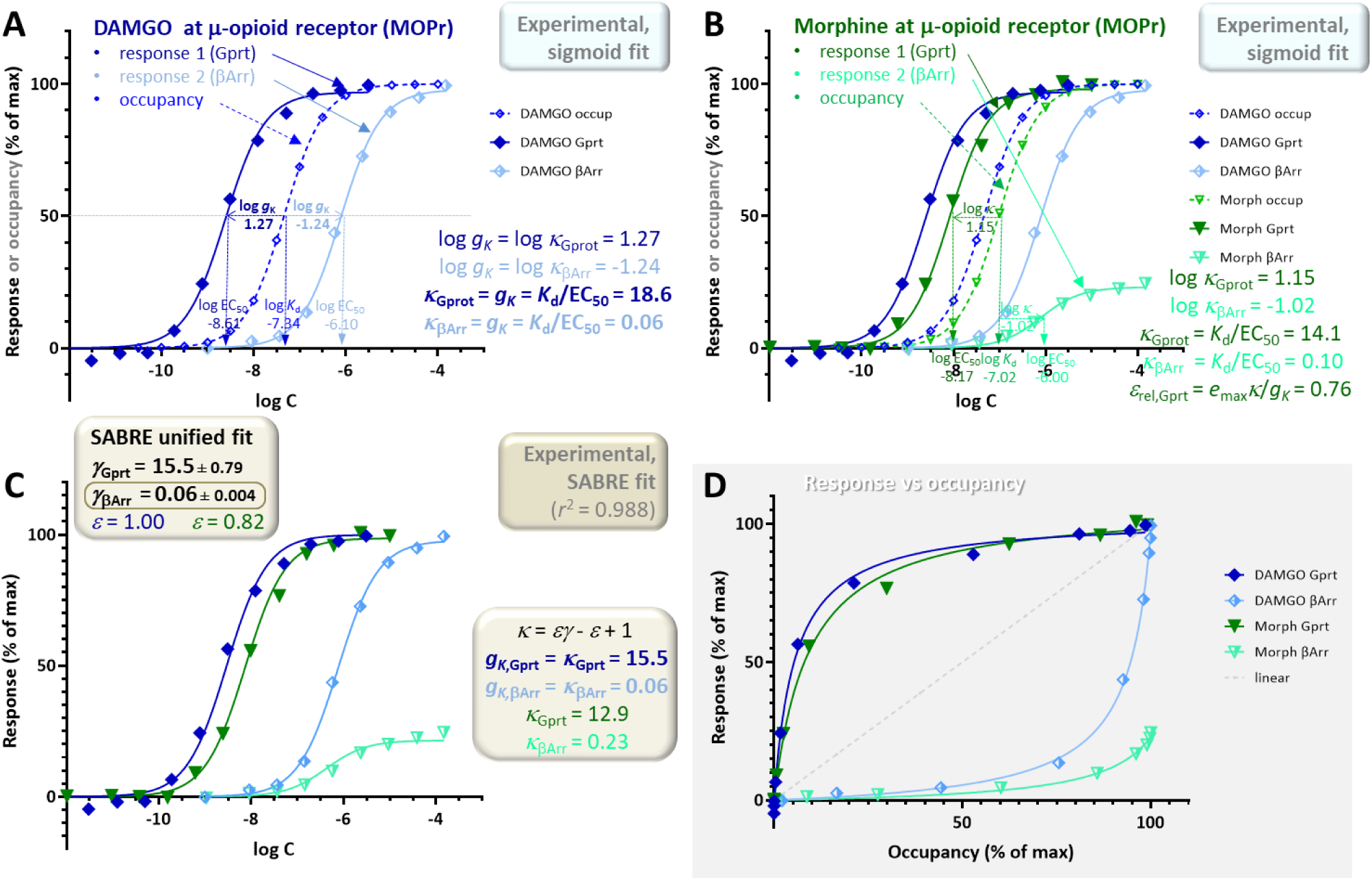
Concentration dependency of receptor occupancy (open symbols and dashed lines) and two different responses measured along different downstream pathways (G_αi2_ activation as well as β-arrestin2 recruitment – darker closed and lighter half-closed symbols, respectively with continuous lines) at the μ-opioid receptor (MOPr) for DAMGO (**A**, blue) and morphine (**B**, green). Experimental data [54] and the corresponding *K*_d_ and EC_50_ estimates were used to obtain the two different *g_K_* gain parameters (eq. 5) (**A**) and *κ* values (eq. 13) (**B**). Note that one response is left- and one is right-shifted compared to the occupancy (EC_50,Gprt_ < *K*_d_ < EC_50,βArr_); accordingly, sigmoid fit of the full agonist DAMGO data indicated gains *g_K_* of 18.6 and 0.06 for the G-protein and β-arrestin responses, respectively (**A**). As before in Figure 10, the same data was also fitted with SABRE (eq. 19) to determine the corresponding gain (*γ*_Gprot_, *γ*_βArr_) and efficacy parameters; however, here the same efficacy was assumed for both pathways (i.e., *ε* = *ε*_Gprot_ = *ε*_βArr_; no bias) (**C**). The gain parameters obtained from the unified fit of all data by SABRE (15.5 ± 0.8 and 0.06 ± 0.004; **C**) also indicate *γ*_βArr_ < 1, i.e., signal attenuation (dampening) and not amplification in this pathway. Because of this, the relatively weak response in the β-arrestin pathway for morphine can be fitted by SABRE without having to assume biased response (*ε* = *ε*_Gprt_ = *ε*_βArr_ = 0.82 ± 0.02); lack of a clearly biased response is also noticeable in the response versus occupancy (*f*_resp_ vs *f*_occup_) graph (**D**).

As for all other cases, these data can also be fitted within the unified framework of SABRE; however, this requires extending the range of the gain parameter *γ* by allowing it to have values less than one, *γ* < 1, i.e., modeling the response in the corresponding pathway as apparent signal attenuation (dampening) (Figure 11C). With this extension, good fit can be obtained (*r*^2^ = 0.988; Figure 11C) even with a four-parameter version model (*γ*_Gprt_, *γ*_βArr_, *ε*_DAMGO_, *ε*_morphine_), i.e., without having to assume different efficacies for the different pathways (*ε* = *ε*_Gprt_ = *ε*_βArr_; no biased agonism). The gain parameters obtained from SABRE (*γ*_Gprt_ = 15.5 ± 0.8 and *γ*_βArr_ = 0.06 ± 0.004; Figure 11C) are in good agreement with those from the sigmoid fit of the full agonist data (18.6 and 0.06; Figure 11A). Thus, there is consistent evidence for *g_K_*_,βArr_ indicating signal attenuation and not amplification in this pathway. Because of the combination of the less than unity gain in the β-arrestin pathway (*γ*_βArr_ = 0.06 ± 0.004) and ligand efficacy of morphine (*ε*_morphine_ = 0.82 ± 0.02), the relatively weak β-arrestin response of morphine can be fitted without having to assume biased agonism (i.e, *ε*_Gprot_ = *ε*_βArr_). The lack of a biased response is also noticeable in the response versus occupancy graph (Figure 11D), where the responses for morphine and DAMGO follow the same pattern contrary to the case of TRV023 and angiotensin II in the previous example (Figure 10D). This is even more evident in a bias plot, which shows the response produced in one pathway directly as a function of the response produced in the other (i.e., *f*_resp2_ vs *f*_resp1_) (Figure 12B). Thus, these data suggest that morphine produces a relatively weak β-arrestin response due to a combination of its partial agonism at MOPr and the apparent signal attenuation in this pathway and not due to biased agonism (similar to the case of oliceridine mentioned earlier [8, 52]).

**Figure 12.**
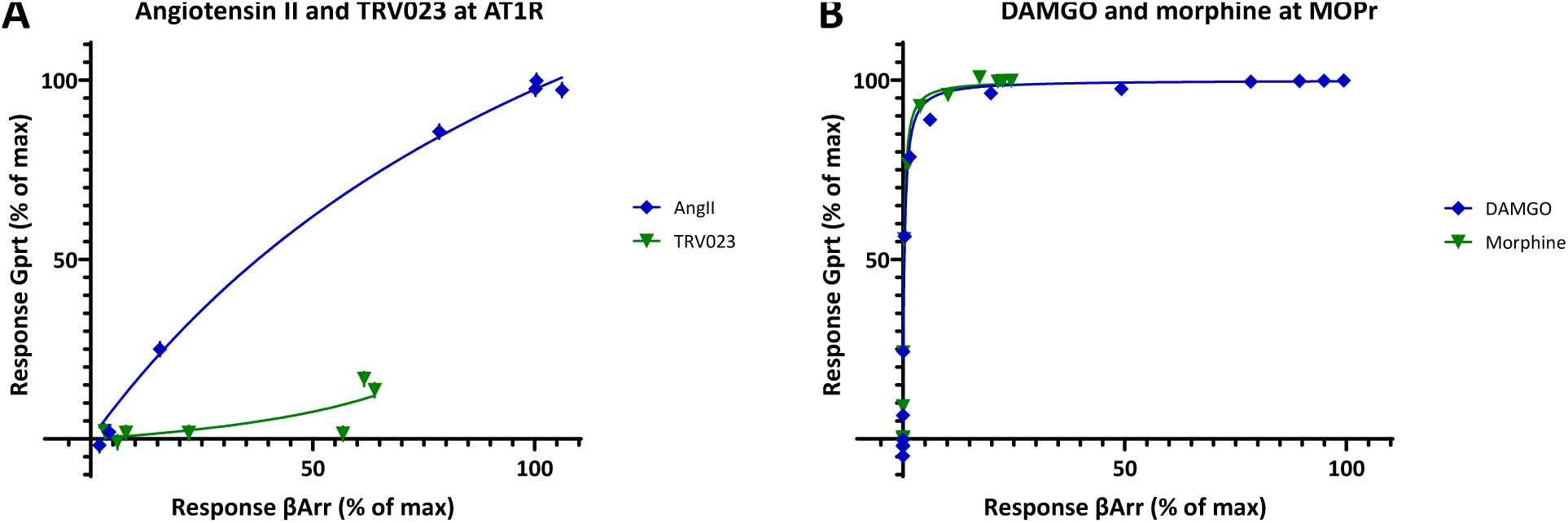
Bias (relative response) plots showing one fractional response as a function of the other (*f*_resp1_ vs *f*_resp2_) for the angiotensin II receptor 1 (AT1R) data from Figure 10 (**A**) and the μ-opioid receptor (MOPr) data from Figure 11 (**B**). While a β-arrestin–biased response for TRV023 is evident when compared to the balanced (nonbiased) agonist AngII (**A**), there is no clear indication of bias for morphine as compared to DAMGO (**B**). However, indications of bias could be somewhat masked in **B** as the plots are strongly curved due to the large difference between the amplifications in these pathways (>250-fold). AT1R data could be used directly as it was (**A**) as both responses were measured at the same agonist concentrations; for the MOPr data (**B**), interpolated values had to be used for the β-arrestin response as they were not measured at the same concentrations as the G-protein response.

### Spare Receptors – a Misnomer

As discussed here, in many systems, maximal or close to maximal response can be achieved when only a relatively small fraction of the receptors is occupied. In current pharmacological terminology, such systems are said to possess “*spare receptors*” or “*receptor reserve*”. According to IUPHAR *spare receptors* are assumed to exist if “a full agonist can cause a maximum response when occupying only a fraction of the total receptor population” (i.e., *f*_resp_ ≈ 1 with *f*_occup_<1) [31]. Similarly, in widely used textbooks, s*pare receptors* are said to be present if “it is possible to elicit a maximal biologic response at a concentration of agonist that does not result in occupancy of all of the available receptors” (*Katzung’s Basic and Clinical Pharmacology* [7]) or “*spare receptors* are said to exist if the maximal drug response (*E*_max_) is obtained at less than 100% occupation of the receptors (*B*_max_)” (*Katzung & Trevor’s Pharmacology Examination & Board Review* [57]). They are judged to be “present if *K*_d_ > EC_50_”, and “in practice, the determination is usually made by comparing the concentration for 50% of maximal effect (EC_50_) with the concentration for 50% of maximal binding (*K*_d_). If the EC_50_ is less than the *K*_d_, *spare receptors* are said to exist” [57] (Figure 13A). These criteria are, however, very much in line with the presence of signal amplification and the definition of the gain factor *gκ* as introduced here (eq. 5) since EC_50_ < *K*_d_ implies *gκ* = *K*_d_/EC_50_ > 1, i.e., amplification. *Spare receptor* implies a “receptor that does not bind drug when the drug concentration is sufficient to produce maximal effect” [57]. Considering the standard definition of spare, i.e., “additional to what is required for ordinary use”, this suggests that they are only used in special circumstances the way, e.g., spare tires or spare rooms are, but this is not the case. In these systems, there is indeed an excess or surplus of receptors, as receptors do not need to be all occupied to achieve maximal response; however, there is no special population of “spare” receptors in addition to that of “regular” receptors that start to fill up only after the “regular” ones are all occupied, as their name might imply. As more correctly described in another textbook (*Rang and Dale’s Pharmacology* [6]): “The existence of spare receptors does not imply any functional subdivision of the receptor pool, but merely that the pool is larger than the number needed to evoke a full response”. Accordingly, *spare receptor*, which is a misnomer, should be avoided. If needed, *receptor reserve*, which is less frequently used, is a better, less confusing terminology [31]. In some cases, it is feasible that there is a *receptor reserve* in the sense they are in excess and do not contribute to the maximum response because of limited downstream transducer availability, so that even if occupied, they cannot generate a response (e.g., only half of GPCRs have access to a G-protein). However, these receptors are again not part of any special *spare* population, and they are subject to being occupied by a ligand just as all others even if they might not be able to initiate signal transduction resulting in a response.

**Figure 13.**
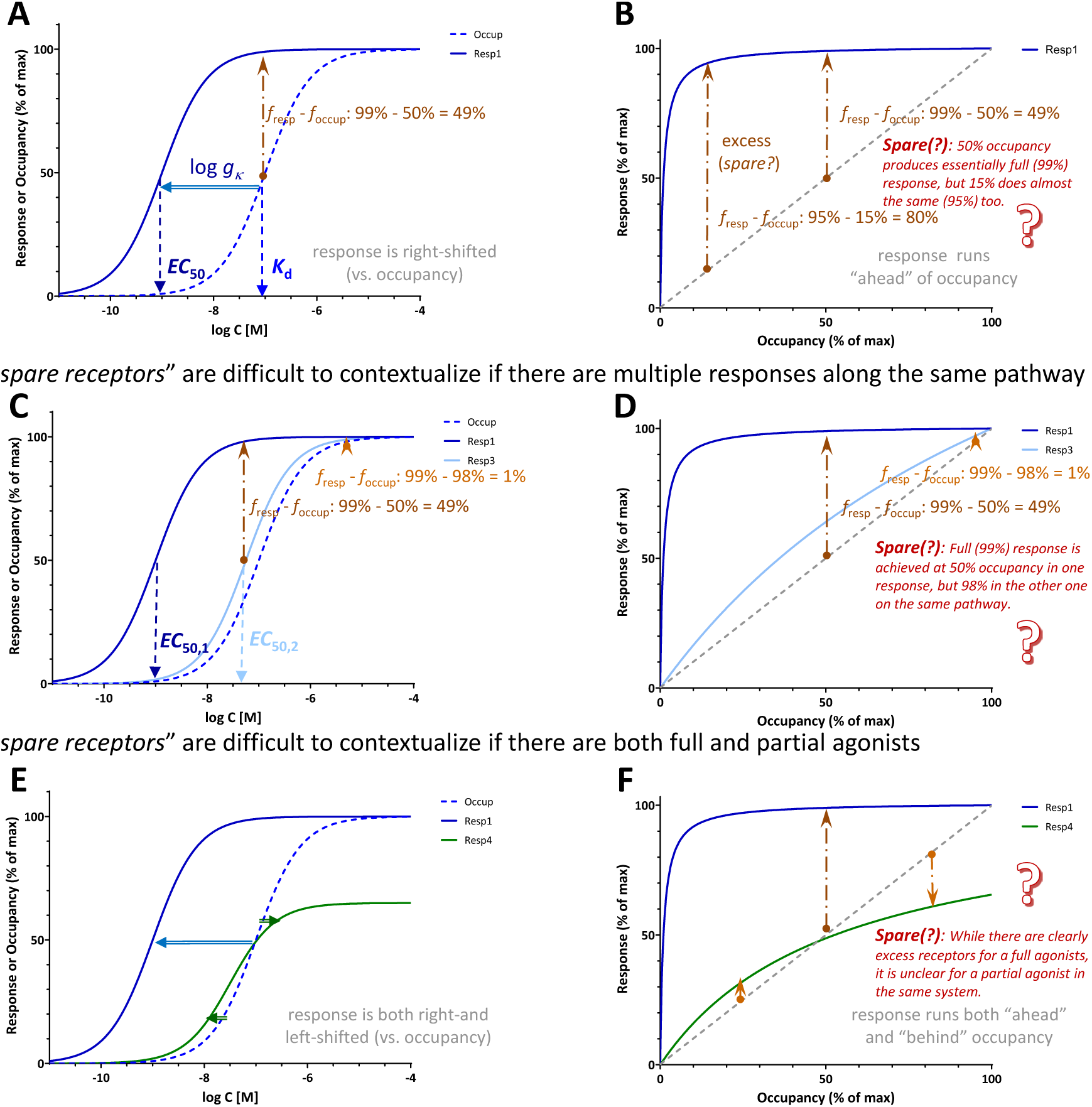
Difficulties related to the quantification of *spare receptors*. They are assumed to exist if (essentially) maximal response is obtained at less than full occupation of receptors and are judged to be present if *K*_d_ > EC_50_ [7, 57]. The excess fraction of receptors as assessed on the vertical axis when comparing response vs occupancy depends on the point of assessment (**A**-**B**); for example, as shown here, the excess is 80% at 95% response but only 49% at 99% response, both of which could be considered essentially maximal responses. They are also difficult to contextualize if two different responses are assessed downstream from the same receptor (**C**-**D**). Here, for example, 99% response is already achieved at 50% occupancy for one response, but only at 98% occupancy for the other even if they can be responses on the same pathway just assessed at different vantage points. *Spare receptors* are also difficult to contextualize if both full and partial agonists are present, as for partial agonists, the response can run both ahead and behind the receptor occupancy (**E**-**F**).

Furthermore, not only is the *spare receptor* term a misnomer, but there is no clear-cut way to quantify them (Figure 13). The percent of excess receptors (i.e., percent response over percent occupancy as judged from the vertical axis) depends on the point of assessment, and if the response is sufficiently left-shifted compared to occupancy, essentially maximal response (e.g., 95% or 99%) can be achieved at (very) different occupancies as illustrated in Figure 13A-B. For the response shown there, 95% of maximum is achieved at 15% occupancy (80% excess) while 99% of maximum at 50% occupancy (49% excess), and these are both essentially maximum responses, typically well within the margin of experimental error. The amount of *spare receptors* is impossible to quantify unequivocally in cases where multiple responses are assessed downstream from the same receptors, such as it was, for example, the case of very differently shifted responses shown in Figure 6. For a corresponding illustration shown in Figure 13C-D, maximal (99%) response is achieved at 50% occupancy in one response, but only 98% occupancy in the other one despite both being on the same pathway downstream from the same receptor just subject to different amplifications. *Spare receptors* are also difficult to contextualize if both full and partial agonists are present because for partial agonists, the response can run both ahead and behind the receptor occupancy. In such cases, it might appear that there are spare receptors when assessed at a low occupancy, but they “disappear” at higher occupancies (Figure 13E-F). Thus, the amount of spare receptors depends, among others, on what is considered essentially maximal response, what response is assessed, and what ligand is used.

Another more complex approach used to demonstrate the existence of *spare receptors* is “by using irreversible antagonists to prevent binding of agonist to a proportion of available receptors and showing that high concentrations of agonist can still produce an undiminished maximal response” [7]. Further, “higher concentrations of antagonist … reduce the number of available receptors to the point that maximal response is diminished”. This is, of course, the classic Furchgott method of irreversible receptor inactivation [15, 16] discussed earlier (example 3; Figure 7). As shown there, the same change in response curves can be produced with the assumption of signal amplification.

Thus, *spare receptor* is a misnomer and a terminology that should be avoided – there is no special pool of “spare” receptors (additional to what is required for ordinary use), just there are excess receptors and not all need to be occupied to elicit full response. In fact, this can be considered a hallmark of a well-engineered system designed to provide some redundancy. Signal amplification with gain as defined here (*gκ* = *K*_d_/EC_50_; eq. 5) is a convenient alternative, and the criteria for its presence, *gκ* = *K*_d_/EC_50_ > 1.0, fully agrees with the textbook definition of the evidence for the presence of *spare receptors*: “EC_50_ is less than the *K*_d_” [57]. Furthermore, while the vertical shift for “excess” varies, the horizontal shift that on semi-log graph corresponds to the log gain, log *gκ* = log *K*_d_ – log EC_50_, remains constant (for full agonists) regardless of the point of assessment; thus it is also intuitive and easy to visualize (Figure 4, Figure 13A-B). As discussed, there are many cases where such signal amplification is needed to achieve sufficient sensitivity, and there are many cases where they have physiological or therapeutic relevance [23, 25, 29, 58, 59].

## Conclusions

In conclusion, a signal amplification-based approach can account for complex receptor-occupancy curves where responses run both ahead and behind fractional occupancy (Figure 5), for differences caused in the response by altering receptor levels (e.g., by partial irreversible inactivation such as in the Furchgott method; Figure 8), as well as for different responses at different readout points either downstream on the same signaling pathway (Figure 6) or along diverging pathways caused by balanced and biased agonists (Figure 10, Figure 11). Signal amplification downstream from receptors can be conveniently quantified using the gain parameter *g_K_* = *K*_d_/EC_50_ measured for full agonists. This gain parameter is analogous to those used elsewhere (e.g., in electronics, *g_V_* = *V*_out_/*V*_in_), and it also corresponds to the *γ* gain parameter of SABRE. Further, it is also an intuitive parameter as on customarily used semi-log representations, it equals the horizontal shift between the response and occupancy curves, log *g_K_* = log *K*_d_ – log EC_50_. The presence of such shift (i.e., *K*_d_>EC_50_) was generally considered as evidence for the existence of *receptor reserve* or *spare receptors*, a misnomer that should be avoided. For partial agonists, the *κ* = *K*_d_/EC_50_ shift is smaller than for full agonists as not all occupied receptors are active, and relative efficacies can be estimated by comparing *e*_max_·*κ* products.

## Supplementary Information

Supplementary information includes Appendix 1 with derivation of the horizontal shift between two sigmoid functions.

## Author Contributions

PB is the sole author; he developed the concept, performed the calculations and data fittings, and wrote the manuscript.

## Acknowledgment

N/A

## Data Availability Statement

Data used for illustrations of model fit are either simulated data generated as described or reproduced from previous publications as indicated in the corresponding figures. The datasets generated and/or analyzed are available from the corresponding author upon reasonable requests.

## Supplementary Information

## Appendices

### Appendix 1. Horizontal shift between two sigmoid functions

Assuming straightforward hyperbolic functions (sigmoidal on the semi-log scale) both for the occupancy and the response, the corresponding fractional occupancies are:

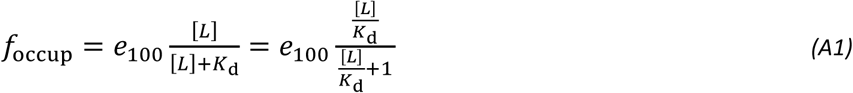

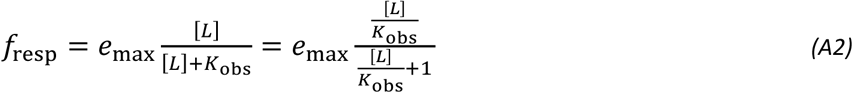

Here, *e*_max_ and *e*_100_ are the corresponding maxima for response and occupancy, respectively (with the assumption that occupancy always reaches its full maximum, hence the notation of *e*_100_ for 100%), and *K*_obs_ and *K*_d_ the half-maximal concentrations. To obtain the ligand concentration values that produce the same (fractional) response and occupancy [L]_o_ and [L]_r_, respectively, one can set *f*_occup_ = *f*_resp_ (any value as long as they are the same):

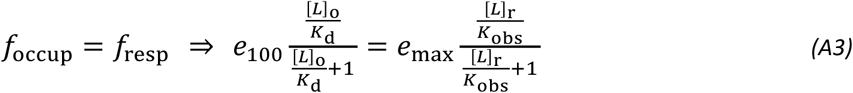

For a full agonist, response also reaches its full maximum (just as occupancy does), *e*_max_ = *e*_100_ = 1 (100%), so that *f*_occup_ and *f*_resp_ follow the same functional form (eqs. A1 & A2) just with a shifted argument. Thus, from eq. A3:

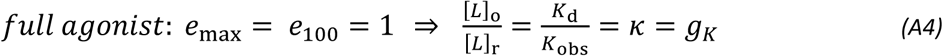

Thus, for a full agonist, the *K*_d_/*K*_obs_ (i.e., *K*_d_/EC_50_) ratio corresponds to the *g_K_* gain parameter as defined earlier, and on the typical semi-log plot, it corresponds to the horizontal shift between occupancy and response, which is the same along any horizontal line:

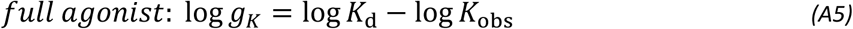

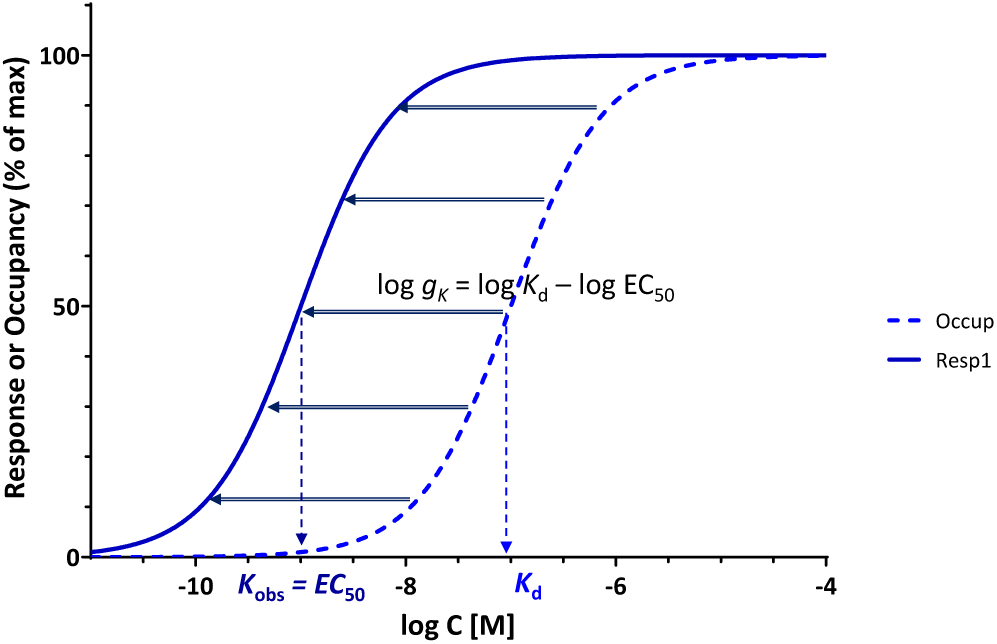

## Footnotes

1 It is notable from a viewpoint comparing the development of the different science fields and a comparison of the sophistication of the theoretical approaches used at a given time that this mathematically relatively simple Clark equation (eq. 2) widely used in pharmacology [9] was published the very same year (1926) as the beautifully complex Schrödinger equation that is in many ways the culmination of quantum mechanics [10]. While the Schrödinger equation may appear simple in its elegant time-dependent form involving the Hamiltonian operator, *H*, and the state vector |*Ψ*(*t*)〉: 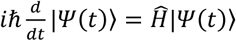 it is in fact a quite complex partial differential equation that governs the wave function of quantum-mechanical systems, as it might be more apparent from its one-dimensional form: 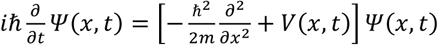

